# Dynamic corticothalamic modulation of the somatosensory thalamocortical circuit during wakefulness

**DOI:** 10.1101/2023.07.18.549491

**Authors:** Elaida D. Dimwamwa, Aurélie Pala, Vivek Chundru, Nathaniel C. Wright, Garrett B. Stanley

## Abstract

The feedback projections from cortical layer 6 (L6CT) to sensory thalamus have long been implicated in playing a primary role in gating sensory signaling but remain poorly understood. To causally elucidate the full range of effects of these projections, we targeted silicon probe recordings to the whisker thalamocortical circuit of awake mice selectively expressing Channelrhodopsin-2 in L6CT neurons. Through optogenetic manipulation of L6CT neurons, multi-site electrophysiological recordings, and modeling of L6CT circuitry, we establish L6CT neurons as dynamic modulators of ongoing spiking in the ventro-posterior-medial nucleus of thalamus (VPm), either suppressing or enhancing VPm spiking depending on L6CT neurons’ firing rate and synchrony. Differential effects across the cortical excitatory and inhibitory sub-populations point to an overall influence of L6CT feedback on cortical excitability that could have profound implications for regulating sensory signaling across a range of ethologically relevant conditions.

## Introduction

Amid the traditionally studied feedforward neuronal pathways underlying sensory perception are numerous feedback processes that are much less well understood. Layer 6 corticothalamic (L6CT) feedback neurons of all primary sensory cortices except the olfactory cortex are one such feedback process. These highly convergent, glutamatergic cortical neurons contribute ∼30-50% of the synapses in sensory first-order (FO) thalamic nuclei (Liu et al., 1995; Sherman & Guillery, 2002; Sherman & Koch, 1986a; Van Horn et al., 2000). Thus, L6CT neurons provide a potent modulatory input on thalamic excitability that likely plays a critical yet enigmatic role in thalamocortical sensory signaling.

In addition to providing direct, monosynaptic excitation to the FO thalamus, L6CT neurons also provide disynaptic inhibition via axon collaterals targeted to the inhibitory thalamic reticular nucleus (TRN), which, in turn, projects to the FO thalamus. Studies in thalamocortical brain slice have shown that the L6CT-mediated confluence of monosynaptic excitation and disynaptic inhibition in the FO thalamus enables a rich set of potential effects on thalamic excitability (Crandall et al., 2015; McCormick & von Krosigk, 1992). While several recent in-vivo studies have attributed L6CT neurons to either enhancing *or* suppressing the gain of FO thalamic sensory responses (Born et al., 2021; Denman & Contreras, 2015; Guo et al., 2017; Mease et al., 2014; Olsen et al., 2012; Pauzin & Krieger, 2018; Spacek et al., 2022; Temereanca & Simons, 2004), the full dynamic range of L6CT modulation on FO thalamus is poorly understood during wakefulness (Guo et al., 2017; Kirchgessner et al., 2020). Moreover, cortical neurons integrate the modulated thalamic inputs as well as direct modulatory effects from intracortical L6CT projections. The effect of the full range of L6CT modulation on cortical neuron activity during wakefulness remains unknown (but see Olsen et al., 2012).

Starting as early as the pioneering neuroanatomical work of Ramón y Cajal (Ramón Y Cajal, 1906), a plethora of studies have outlined the intricate anatomical details of the L6CT circuit (Cudeiro & Sillito, 2006; Guillery & Sherman, 2002; Usrey & Sherman, 2019; Varela, 2014). However, it is only from recent work that we are beginning to understand the precise contexts in which these neurons become engaged and exert their influence, expanding the range of observed L6CT activity patterns. In addition to being responsive to sensory stimuli (Augustinaite & Kuhn, 2020; Briggs & Usrey, 2009; Swadlow, 1989; Zhou et al., 2010), L6CT neuron activity is also modulated by extra-sensory features, such as head rotation (Vélez-Fort et al., 2018), motor-preparation (Clayton et al., 2021), whisking (Dash et al., 2022) and locomotion (Augustinaite & Kuhn, 2020). Additionally, L6CT neurons serve as a circuit mechanism for brain state-dependent modulation of the FO thalamus (Reinhold et al., 2023). Collectively, these findings point to a continuous and context-dependent feedback modulation of the FO thalamus that acts through the regulation of L6CT population activity across a range of behaviors. Yet a unified understanding of the direct effects of this broadly acting regulation mechanism on thalamocortical structures remains elusive.

To causally uncover the full range of influence that L6CT neurons have on ongoing and stimulus-evoked spiking activity throughout the sensory thalamocortical circuit during wakefulness, we targeted silicon probe recordings to the whisker region of the primary somatosensory cortex (S1), the ventro-posterior-medial nucleus (VPm), and the thalamic reticular nucleus (TRN) of awake, NTSR1-cre mice selectively expressing Channelrhodopsin-2 (ChR2) in L6CT neurons. We found that increasing optogenetic activation of L6CT neurons shifted from suppressing to enhancing VPm ongoing activity, which we refer to as bidirectional modulation of VPm. The findings here suggest that L6CT-mediated bidirectional modulation of VPm activity is likely a combination of indirect contributions from L6CT activation of TRN and direct contributions to VPm through changes in L6CT population firing rate as well as spiking synchrony. Differential effects of L6CT activation on ongoing regular-spiking (RS) and fast-spiking (FS) cortical neurons’ activity result in an increase in overall cortical excitability. Further, we found that L6CT neuron activation enhances the magnitude of the VPm sensory response and leaves the sensory response magnitude of the RS S1 population unchanged. Yet, the sensory response magnitude of the FS S1 population is maximally enhanced at lower but not higher optogenetic drive of L6CT neurons, reminiscent of the dynamic changes observed in the ongoing activity of VPm neurons that would then be inherited by S1, resulting in the differential L6CT engagement of cortical excitatory and inhibitory subpopulations. These results highlight L6CT neurons as rate and synchrony-dependent, dynamic modulators of cortical and thalamic excitability.

## Materials and Methods

### Experimental model and subject details

All experiments were conducted in 11 male and 7 female NTSR1-cre (B6.FVB(Cg)-Tg(Ntsr1-cre)GN220Gsat/Mmcd, MMRRC) adult mice as well as 1 wildtype, C57BL/6J mouse, all aged 6 weeks to 6 months housed under a reversed light-dark cycle. All procedures were approved by the Institutional Animal Care and Use Committee at the Georgia Institute of Technology and were in agreement with guidelines e**s**tablished by the National Institutes of Health.

### Headpost implantation

The mice were kept under isoflurane anesthesia (5% vaporized in O2 for induction, 1-1.5% for maintenance), their temperature maintained at ∼37 C, and ophthalmic ointment (Puralube, Fisher Scientific) was applied to the eyes to prevent drying. A stereotaxic frame maintained the animal’s head level in all three planes while a custom stainless steel headpost and recording chamber was affixed to exposed skull with dental cement (C & B Metabond Parkell, Inc). The mice were then provided at least three days to recover (Bolus et al., 2021; Borden et al., 2022; Guo et al., 2014; Pala & Stanley, 2022; Wright et al., 2021, 2021).

### rAAV injection

An image of superficial vasculature was overlaid with a map of the whisker columns of the primary somatosensory cortex (S1), generated functionally via intrinsic signal optical imaging performed through a thinned skull under 1-1.5% isoflurane anesthesia (Liew et al., 2021; Masino et al., 1993; Pala & Stanley, 2022). On a subsequent day, also under 1-1.5% isoflurane anesthesia, 800 nL of the cre-dependent AAV5-EF1a-DIO-hChR2(H134R)-eYFP.WPRE.hGH (UPenn Vector Core; Kirchgessner et al., 2020; Pauzin & Krieger, 2018) was injected at a depth of 800 um in the C1 whisker column. The exposed skull was covered with a silicone elastomer (Kwik-Cast, WPI) and the mice were returned to their home cage and provided at least three days to recover. Electrophysiological recordings were conducted no sooner than 3 weeks after the injection to allow for stable opsin expression.

### Habituation to head-fixation

Mice were habituated to head-fixation for a minimum of 5 days. Head-fixation sessions started with a duration of 15 minutes and went up to 90 minutes. Sessions gradually increased in duration from day-to-day as did the exposure to experimental conditions (e.g., whisker stimulation, exposure to optogenetic stimuli, and paw restraint). Throughout each session, mice were periodically rewarded with sweetened condensed milk (Bolus et al., 2021; Borden et al., 2022; Guo et al., 2014; Pala & Stanley, 2022; Wright et al., 2021).

### Electrophysiological recordings

All electrophysiological recording sessions were conducted in darkness using silicon probes in awake, head-fixed mice periodically rewarded with sweetened condensed milk (Pala & Stanley, 2022). Often, sessions consisted of simultaneous recording from two brain regions. Additionally, sessions were conducted no sooner than two hours after craniotomy procedures under 1-1.5% isoflurane anesthesia where the dura was left intact. No more than four craniotomies were made per animal.

The location of craniotomies to target S1 was based on the map generated via intrinsic signal optical imaging (see above). Recordings of the activity of S1 neurons were conducted using 32 or 64 channel silicon probes (A1×32-5 mm-25-177-A32, A1×32-Poly3-5mm-25s-177-A32, A1×64-Poly2-6mm-23s-160, or A1×64-Edge-6mm-20-177-A64, Neuronexus, Ann Arbor, MI) inserted to depths between 1250 and 1400 um at an angle of 35 degrees from vertical (Pala & Stanley, 2022).

Craniotomies to target neurons in the ventro-posterior-medial (VPm) nucleus of the thalamus were made 2 mm posterior and 2 mm lateral from bregma. Often, a brief electrophysiological recording was first conducted under isoflurane anesthesia to verify the craniotomy location, or to extend the craniotomy as necessary. Recordings of VPm neurons were conducted using 32 channel silicon probes (A1×32-Poly3-5mm-25s-177-A32, Neuronexus, Ann Arbor, MI) inserted in the vertical plane to depths between 3200 and 3600 um (Bolus et al., 2021; Borden et al., 2022; Liew et al., 2021; Wright et al., 2021).

Craniotomies to target neurons of the thalamic reticular nucleus (TRN) were made 2.4 mm posterior, with a medial edge between 1.5 and 2.5 mm from bregma (Lewis et al., 2015). Often, a brief electrophysiological recording was first conducted under isoflurane anesthesia to locate TRN neurons. Recordings of TRN neurons were conducted using 32 channel silicon probes (Buzsaki32 or A1×32-Poly3-5mm-25s-177-A32, Neuronexus, Ann Arbor, MI) inserted in the vertical plane to depths between 2900 and 3200 um (Lewis et al., 2015; Shosaku et al., 1984).

Continuous signals were acquired using either the Cerebus acquisition system (Blackrock Neurotech, Salt Lake City, Utah) or TDT RZ2 Bioamp Processor (Tucker Davis Technologies, Alachua, FL). Signals were amplified, filtered between 0.3 Hz and 7.5 kHz, and digitized at either 30,000 Hz or 24,414.0625 Hz. For each animal, we conducted one recording session per day and up to three recording sessions per craniotomy.

### Optogenetic stimulation

Optogenetic inputs were delivered using a 470 nm light emitting diode (LED, ThorLabs M470F3) coupled to a 400 um diameter optical fiber (ThorLabs M28L01) manually positioned on the skull surface near the recording probe. Command voltages for optogenetic inputs were controlled by custom scripts in MATLAB and Simulink Real-Time (MathWorks, Natick, MA), sampled at 1 kHz.

All optogenetic input began with a ramp of the LED intensity that was 250 ms in duration, serving to minimize transient neuronal effects from rapid increases in LED intensity as well as to minimize animal whisking (Urbain et al., 2015). In ramp-and-hold LED trials, the LED intensity was then held fixed at values ranging from 4 to 30 mW/mm^2^ for 500-650 ms. Manipulations to vary L6CT synchrony included LED inputs that were 500-650 ms in duration labelled as either 500 Hz, 100 Hz, or 10 Hz trials. The input for 500 Hz trials consisted of uniformly distributed white noise. Inputs for the 100 Hz and 10 Hz trials consisted of uniformly distributed white noise that was low-pass filtered (3^rd^ order Butterworth filter) at their respective frequencies. For all trials, the inter-trial interval was a minimum of 3.5 times the duration of the LED input with added uniformly distributed random jitter per-trial ranging from 0 - 0.1/0.5 ms.

### Whisker stimulation

For sensory stimulation, an individual whisker contralateral to the electrophysiological recording site was threaded into a narrow, 1.5 cm long tube connected to a precise, computer-controlled galvanometer (Cambridge Technologies, Worthington, MN; sampled at 1 kHz; Borden, Liew, Pala, Wright). The whiskers were deflected at a velocity of 300 deg/s in the caudo-rostral direction with a “sawtooth” waveform consisting of rise and fall times of 8 ms, where velocity is defined by the average velocity over the rising phase. In trials with concurrent LED input and sensory stimulation, the sensory stimulus was presented 750 ms after the onset of the LED input.

### Videography and whisker motion analysis

The movement of the whiskers ipsilateral to the electrophysiological recording site was captured under infrared illumination using CCD cameras (DMK 21BU04.H USB 2.0 monochrome industrial camera, The Imaging Source, LLC, Charlotte, NC; or EoSens CL MC1362, Mikrotron, Germany). Frames were individually triggered at 30 Hz or 200 Hz. Using FaceMap (Syeda et al., 2022), whisking “energy” was extracted from manually defined regions of interest (ROIs) around the face. Using custom scripts, we computed the across-frame variation of the sum squared motion energy across ROIs. This time series was then smoothed and time points above a fixed threshold were classified as whisker motion epochs (Pala & Stanley, 2022; Wright et al., 2021). To measure the specific effects of our manipulations unconfounded by effects associated with whisking, all trials included for analysis of ongoing activity had no whisker motion occurring during the trial. For the analysis of sensory responses, only trials in which there was no whisker motion for 75 ms before and after the stimulus were included.

### Histology

For histological validation of the recording location, during one of the electrophysiological recording sessions per craniotomy, the probe was coated with DiI (0.2 mg/mL in ethanol, Invitrogen) to enable fluorescence imaging of the probe track (Liew et al., 2021; Pala & Stanley, 2022). In brief, after the final recording, mice were transcardially perfused with 1x PBS (137 mM NaCl, 2.7 mM KCl, and 10 mM PB, VWR) and then 4% PFA. The brains were extracted, post-fixed overnight in the PFA solution, and then sliced to 100 um thick sections on a vibratome (VT1000S, Leica Biosystems Deer Park, IL). The slices were then incubated with DAPI (2 mM in PBS, AppliChem, Council Bluffs, Iowa) for 15 min, mounted on slides with a DABCO solution (1,4-Diazabicyclo[2.2.2]octane, Sigma), and imaged using a confocal microscope (Laser Scanning Microscope 900, Zeiss, Germany). Corresponding brain atlas sections were chosen manually (Franklin & Paxinos, 3^rd^ edition).

### Spike sorting and identification of single neuron clusters

Individual recording sweeps were high-pass filtered (3^rd^ order Butterworth filter; 500Hz cut-off frequency), median-filtered across channels, and concatenated before being passed through Kilosort2 for automated spike sorting based on template-matching (Pachitariu et al., 2023). The data were then manually curated in Phy2 (Rossant & Harris, 2013). With the exception of the control experiments conducted in a wildtype mouse, all neuron clusters considered for analysis passed two criteria: a signal-to-noise ratio of the mean spike waveform greater than three and less than 2% of all spikes violating a 2 ms imposed refractory period. Note that the use of more strict inclusion criteria does not change the nature of any claims presented here.

### Identification and classification of S1 neurons

All S1 neurons retained for analysis are required to be sensory stimulus-responsive, defined by passing a minimum of two out of three tests assessing if the stimulus caused a significant change in the spiking activity. The tests were conducted on spikes occurring in 50 ms a pre- and post-stimulus window. The first test was the two-sided Wilcoxon signed rank test on the pre- versus post-stimulus spike counts across trials. The second test determined whether zero overlapped with the 95% bootstrap confidence interval from the stimulus-evoked spike counts subtracted by the pre-stimulus spike counts. The final test used a post-stimulus peri-stimulus time histogram (PSTH) with 10 ms bins to determine whether a minimum of three bins were above or two bins were below the 95% bootstrap confidence interval of pre-stimulus activity (Pala & Stanley, 2022).

S1 neurons were then classified as regular-spiking (RS) or fast-spiking (FS) based on the width of the mean spike waveform, defined as the trough to peak time (Barthó et al., 2014). Neurons with a TPT greater than 0.5 ms were classified as an RS neuron and neurons with a TPT less than 0.4 ms were classified as an FS neuron. Neurons with TPT widths between 0.4 ms and 0.5 ms were not analyzed (Barthó et al., 2014; Pala & Stanley, 2022; Sederberg et al., 2019).

Each neuron was then assigned to a layer relative to the center of layer 4, which was identified functionally from the sensory response on each channel of the recording probe. In brief, we extracted the trial-averaged sensory response for each channel across the following measures: threshold-crossings of the high-pass filtered recording trace, the local field potential (LFP), and the sources computed using current-source density analysis (Sederberg et al., 2019; Sofroniew et al., 2015). We extracted the channels with the earliest onset and largest response across these data types and the center of layer 4 was determined based on collective evidence. All channels, and therefore the neurons recorded on those channels, were then assigned to a layer based on their distance from the determined center of layer 4. Each layer was assigned the appropriate number of channels to cover the following thickness: 2/3: 300 um; 4: 200 um; 5: 300 um; 6: 300 um (Hooks et al., 2011; Pala & Stanley, 2022; Sederberg et al., 2019).

### Classification of L6CT neurons

To identify L6CT neurons, we used the stimulus-associated spike latency test (SALT) to statistically determine whether 5 ms, square LED pulses cause a significant increase in each neuron’s spiking with low temporal jitter across trials (Kvitsiani et al., 2013). In brief, SALT was used to compare the distribution of first spike latencies to the LED pulse compared to distributions created from spontaneous activity. Any deep, RS neuron with a p-value less than 0.01 was classified as a L6CT neuron. Note that L6CT neurons were not required to be sensory stimulus-responsive to be retained for analysis.

### Classification of VPm and TRN neurons

Neurons classified as VPm or TRN neurons are sensory stimulus-responsive neurons recorded at the relevant stereotaxic coordinate for each location. To determine whether a neuron is stimulus-responsive, we individually deflected multiple whiskers with a 10 Hz adapting train of sawtooth, punctate stimuli for one second. We extracted each neuron’s response latency and magnitude to the first and last stimulus in the train in a 30 ms quantification window. VPm and TRN neurons were required to be significantly responsive to both the first and last stimulus in the train for any deflected whisker. Significance was defined using the two-sided Wilcoxon signed rank test to determine if the sensory stimulus caused a significant increase in the spike counts across trials compared to pre-stimulus (spontaneous) spike counts in an equivalent window. The neurons were further required to have an average first spike latency of less than 15 ms for the first stimulus and less than 20 ms for the last stimulus in the train to ensure the recorded neurons were not of the posteromedial nucleus. Finally, each stimulus-responsive neuron was assigned a primary whisker based on the whisker that evoked the earliest and largest response to the first stimulus in the train (Ahissar et al., 2001; Borden et al., 2022; Liew et al., 2021; Wright et al., 2021).

Previous work has shown that TRN axonal spikes can be recorded in extracellular recordings in the VPm and the mean waveform TPT width of these spikes is more narrow than VPm spikes (Barthó et al., 2014; Guo et al., 2017; Lewis et al., 2015; Makinson et al., 2017). Thus, a final inclusion criterion for VPm neurons was that they must have a waveform width TPT of greater than 0.3 ms.

### Measures of neuronal activity

#### Change in ongoing firing rate due to the LED stimuli

It was necessary to quantify the properties of the neuronal circuit when L6CT neuron activity had reached a steady state from optogenetic manipulation. Thus, we omitted the early, more transient modulatory dynamics from our quantifications and defined the effect as the average firing rate across trials in the window from 250 ms (the end of the LED ramp) to 750 ms after the onset of the LED input. These firing rates were subtracted on a trial-by-trial basis from the pre-LED firing rate (spontaneous activity) in an equivalent 500 ms window.

#### Change in ongoing synchrony

To account for changes in firing rate between conditions, the synchrony strength was determined for pairs of neurons by computing the total numbers of spikes in a +/− 7.5 ms window in the resulting cross-correlogram and normalizing by the total number of spikes from each neuron (N_neur1_ and N_neur2_; Liew et al., 2021; Temereanca et al., 2008):

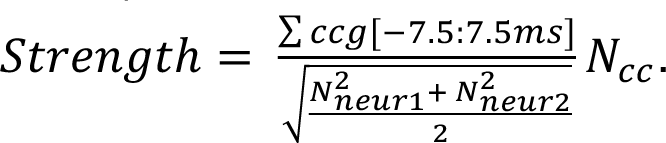

#### Sensory response

Sensory responses were quantified by looking at the average spike rate across trials in a 30 ms window post-sensory stimulus, subtracted on a trial-by-trial basis from the pre-sensory stimulus spike rate in an equivalent 30 ms window. The sensory response for each neuron was determined based on stimulation of its primary whisker.

#### Burst spikes

VPm burst spikes are defined as two or more spikes occurring with an inter-spike interval of less than 4 ms, preceded by 100 ms or more of no spikes, as characteristic of T-type calcium channel burst spikes (Borden et al., 2022; Lu et al., 1992; Swadlow & Gusev, 2001; Whitmire & Stanley, 2016; Wright et al., 2021).

### Statistical analyses

The two-sided Wilcoxon signed rank test with a Bonferroni correction was used between pairs of LED conditions to test for statistically significant changes in various measures of the response of a respective population of neurons. The Spearman’s rank correlation test was used to determine whether increasing LED intensity had a monotonic effect on changes in the ongoing firing rates of the respective population of neurons. When testing for changes in the ongoing activity of individual neurons, the spontaneous versus LED-evoked spike rates were compared using the two-sided Wilcoxon signed rank test. All tests were implemented using MATLAB built-in commands.

### LIF Model

We constructed a simple circuit model that was representative of the key dynamics of the interactions between the L6CT neurons, the VPm, and the TRN. Rather than a largescale network model designed to represent the diversity of properties at the population level, the model we constructed was a reduced circuit model in the spirit of models previously used to describe thalamocortical dynamics (Pinto, 2003), in which the average activity of neuronal populations is captured.

#### Model structure

The model consists of two main nodes, one representing the average VPm neuron response and the other representing the average TRN neuron response, both modelled as leak integrate-and-fire (LIF) neurons. Each node receives noisy excitatory inputs for spontaneous spiking. These inputs are modelled as homogeneous Poisson spike trains, each with their own firing rate. Both nodes also integrate the same 35 excitatory inputs that represent L6CT neuron spiking, modelled as homogenous Poisson spike trains. Finally, the TRN node forms an inhibitory synapse onto the VPm node.

#### Controlling L6CT synchrony

To control the synchrony of the L6CT inputs, the spike times for each of the L6CT spike trains are generated by drawing spikes from two homogeneous Poisson spike trains: one shared and one individual spike train, each with equivalent firing rates. For each synchrony condition, we selected the shared probability, a value between 0 and 1, that determines the likelihood that the spikes for each L6CT input would come from the shared spike train versus the individual spike train. The higher the shared probability, the more likely that the 35 L6CT inputs will have coincident spike times, increasing their synchrony, while maintaining equivalent firing rates.

#### Evolution of model dynamics

Each LIF node’s membrane potential, *V*, evolves according to the membrane equation:

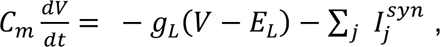

where *C_m_* is the membrane capacitance, *g_L_* and *E_L_* are the leak conductance and reversal potential, respectively. 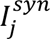 represents the synaptic currents received by each node from the *j^th^*^;^source. Each source is modelled as one synapse according to the equation:

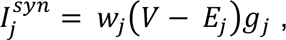

where *w_j_*, *E_j_*, and *P_j_* are the weight, reversal potential, and conductance of the *j^th^*^;^synapse. The synaptic conductance is incremented by *g_jmax_* after each presynaptic spike and decays according to the equation:

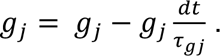

where *τ_gj_* is the respective synapses’ decay constant. To account for synaptic transmission delays, a 1 ms delay is implemented between the time in which a pre-synaptic source fires a spike and the time in which this spike can begin to evolve the current in the post-synaptic neuron.

Finally, we implemented a 2 ms refractory period, during which the evolving currents for the node in a refractory period could not affect the neuron’s membrane potential.

All model parameters were initially based on previously published parameters (Destexhe et al., 1998) and experimental observations (Crandall et al., 2015). The final parameters (Table 1) were then further tuned to better match the experimental neuronal response to the lowest LED input intensity, then allowing the response to all other LED inputs to emerge.

**Table 1.** Parameters used in the LIF model.

| **Table 1. Parameters used in the LIF model** |  |  |
| --- | --- | --- |
| **Variable** | **Value used in model of VPM neuron** | **Value used in model of TRN neuron** |
| $C_m$ | $0.00029 \mu F$ | $0.00014 \mu F$ |
| $g_L$ | $0.029 \mu S$ | $0.007 \mu S$ |
| $E_L$ | $-70 mV$ | $-80 mV$ |
| $w_j$ | $w_{TRN} = 0.075$ | $w_{VPM} = 0$ |
| | $w_{CT} = 1$<br>$w_{Spont.Inp.} = 0.0129$ | $w_{CT} = 1$<br>$w_{Spont.Inp.} = 0.0057$ |
| $E_j$ | $E_{TRN} = -80\text{ mV}$<br>$E_{CT} = 0\text{ mV}$<br>$E_{Spont.Inp.} = 0\text{ mV}$ | $E_{VPm} = 0\text{ mV}$<br>$E_{CT} = 0\text{ mV}$<br>$E_{Spont.Inp.} = 0\text{ mV}$ |
| $\tau_{P,j}$ | $\tau_{P,TRN} = 0.01\text{ s}$<br>$\tau_{g,CT} = 0.1\text{ s}$<br>$\tau_{g,Spont.Inp.} = 0.01\text{ s}$ | $\tau_{P,TRN} = 0.01\text{ s}$<br>$\tau_{g,CT} = 0.1\text{ s}$<br>$\tau_{g,Spont.Inp.} = 0.01\text{ s}$ |
| $g_{jmax}$ | $g_{TRN,max} = 1\text{ }\mu S$<br>$g_{CT,max} = 0.003\text{ }\mu S$<br>$g_{Spont.Inp.,max} = 1\text{ }\mu S$ | $g_{VPm,max} = 1\text{ }\mu S$<br>$g_{CT,max} = 0.003\text{ }\mu S$<br>$g_{Spont.Inp.,max} = 1\text{ }\mu S$ |
| $V_{threshold}$ | $-54\text{ mV}$ | $-54\text{ mV}$ |
| $V_{reset}$ | $-80\text{ mV}$ | $-80\text{ mV}$ |

## Data and code availability

The data from this study along with analysis code can be made available upon request.

## Results

To investigate the direct modulatory effect of L6CT neurons on thalamocortical spiking activity during wakefulness, we injected a recombinant adeno-associated virus (rAAV) encoding Cre-dependent ChR2 fused with eYFP into the whisker region of S1 of NTSR1-Cre mice (Gong et al., 2007). ChR2-expressing L6CT neurons were optically driven with ramp-and-hold LED inputs (250 ms ramp, 500 ms hold durations) across a range of intensities using a 470 nm LED coupled to a 400 um diameter optical fiber positioned at the S1 surface. Concurrently, we targeted silicon probe recordings to S1 and measured the spiking activity of populations of individual L6CT neurons. L6CT neurons were distinguished from other S1 neurons based on a statistical measure of their short latency and low jitter spike activation to a 5 ms, square LED pulse, quantified using the stimulus-associated spike latency (SALT) test (Figure 1b; Kvitsiani et al., 2013). The analysis of neuronal activity both here and in subsequent analysis is during periods of whisker quiescence as determined from videography (See Materials & Methods).

**Figure 1.**
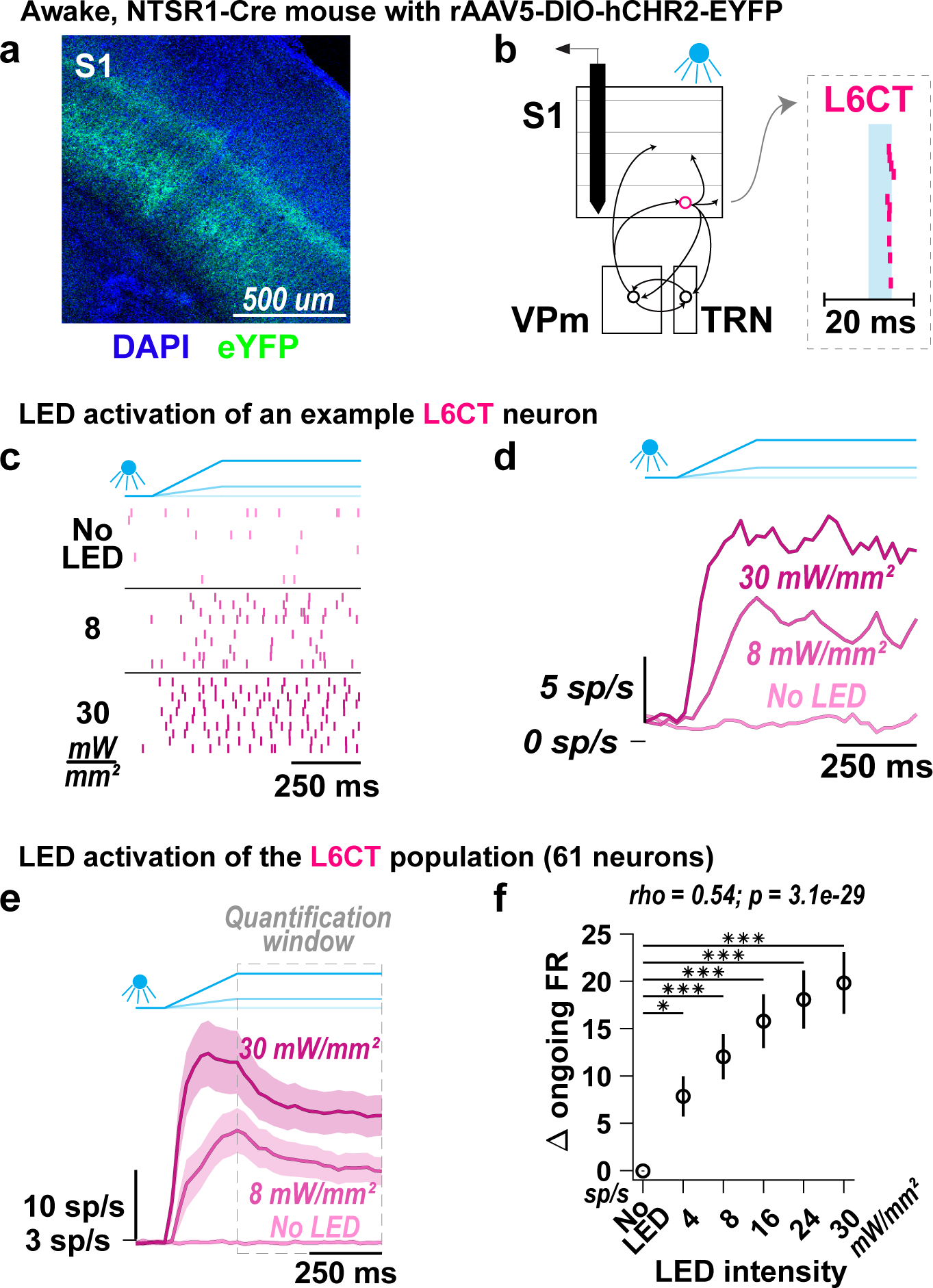
Optogenetic activation of L6CT neurons in awake mice. **a.** A coronal section of DAPI labelled S1 neurons (blue) and selective ChR2-eYFP expression in L6CT neuron prcoesses (green) of NTSR1-Cre mice. **b.** The direct effect of the LED on L6CT neuron activity is measured from in vivo, silicon probe recordings of neuronal spiking in S1 of awake, head-fixed, NTSR1-Cre mice. L6CT neurons are activated by an optical fiber positioned at the cortical surface and are identified by their low latency and low jitter spiking response to 5 ms LED pulses (right inset raster; see Materials and Methods). **c.** Raster plots of an example L6CT neuron. This neuron’s spiking increases to 8 and 30 mW/mm^2^ LED inputs. **d.** PSTHs of the same L6CT neuron in ***c***. **e.** Grand PSTHs (mean +/− sem across neurons) of the L6CT population. The population’s firing rate is enhanced by increasing LED intensity. **f.** The mean +/− sem change in ongoing firing rate of the L6CT population contributing to the PSTHs in ***e*** is monotonically enhanced by increasing LED intensities (8 mW/mm^2^: 12.00 +/− 2.45 sp/s, p = 6.9e-6; 30 mW/mm^2^: 20.80 +/− 3.43 sp/s, p = 9.7e-12; p-value indicates pairwise comparison to No LED condition using the two-sided Wilcoxon signed rank test with a Bonferroni correction; * indicates p < 1.0e-2; *** indicates p < 2.0e-4; rho & p-value: Spearman’s rank correlation).

For the example L6CT neuron presented in Figure 1c, increasing LED intensity increases the neuron’s spiking. A relatively rapid rise in firing rate during the LED ramp followed by steady-state spiking during the hold period of the LED input is apparent in both the individual neuron population peri-stimulus time histograms (PSTHs; Figure 1d) as well as the population grand PSTH (Figure 1e). Here and in subsequent analyses, we quantified the effects of the LED activation during this period of steady-state activity of the L6CT population, as highlighted by the “quantification window” in Figure 1e. Across the population of recorded L6CT neurons, the measured change in ongoing firing rate increases monotonically with LED intensity (Figure 1f). Note that the change is relative to a fairly low baseline level of firing of approximately 2.6 spikes/second of the L6CT neurons (compared to a baseline level of firing of 4.9 spikes/second for the rest of the S1 RS neuron population). Also note that the elicited changes in spiking activity were not due to the LED illumination alone, as verified in control experiments conducted in a wildtype mouse (Figure S1).

### Increasing optogenetic activation of L6CT neurons results in bidirectional modulation of ongoing firing rates in the VPm

We next evaluated the direct effect of L6CT neuron activation on the FO thalamus by targeting silicon probe recordings to the ventro-posterior-medial (VPm) nucleus of the thalamus and measuring the spiking activity of populations of individual VPm neurons during optogenetic activation of L6CT neurons with increasing LED intensity (Figure 2a). The identification of VPm neurons recorded at the relevant stereotaxic coordinates was validated based on measures of their sensory response and spike waveform width (Figure 2b & S2; Ahissar et al., 2000, 2001; Borden et al., 2022; Liew et al., 2021; Wright et al., 2021; see Materials & Methods).

**Figure 2.**
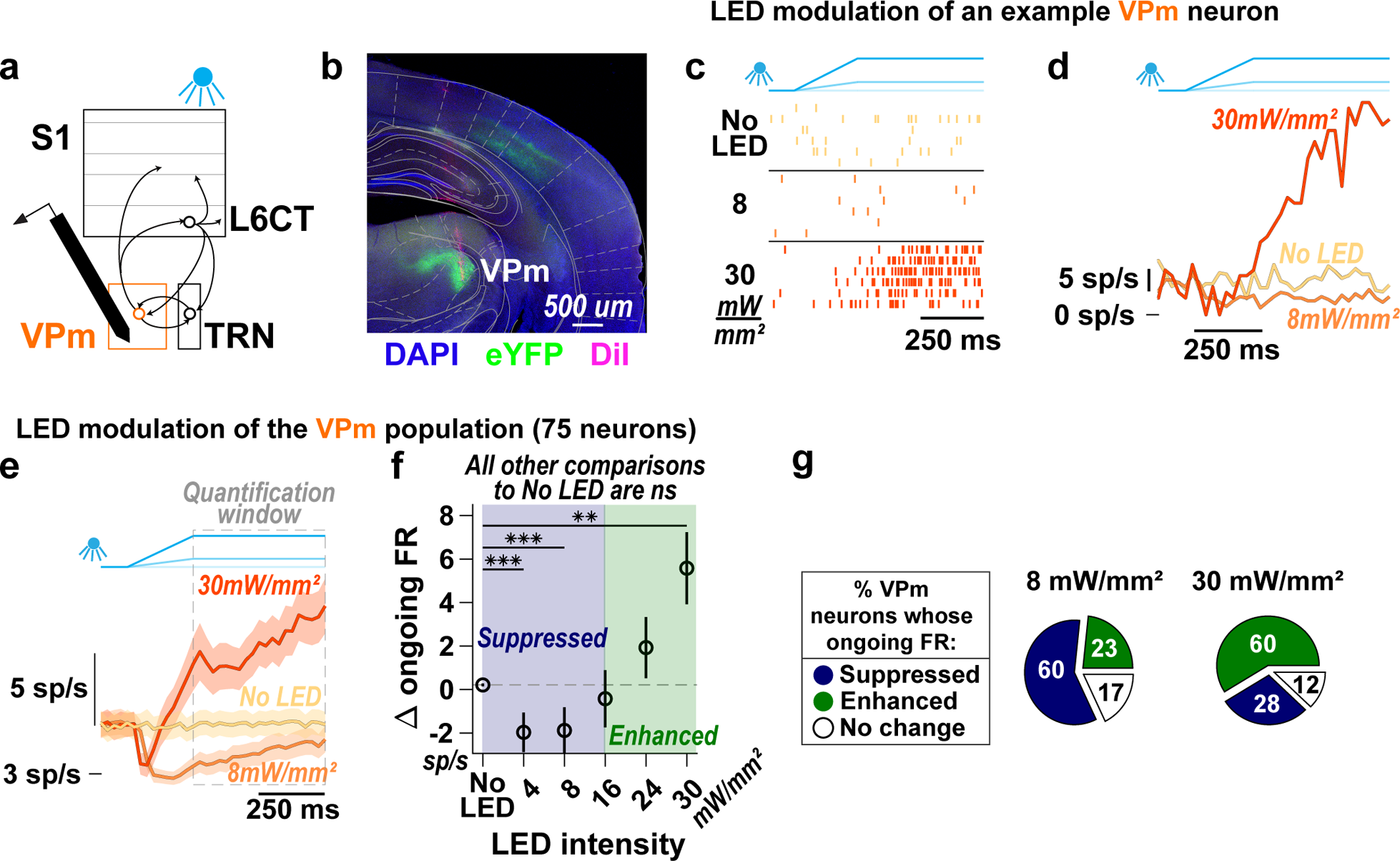
Increasing optogenetic activation of L6CT neurons results in bidirectional modulation of ongoing firing rates in the VPm. **a.** The direct effect of L6CT neuron activation on VPm is measured from in vivo, silicon probe recordings of neuronal spiking in the VPm of awake NTSR1-Cre mice selectively expressing ChR2 in L6CT neurons. **b.** A coronal section of DAPI labelled neurons (blue), ChR2-eYFP expression in L6CT axons terminating in the VPm (green), and a VPm recording tract labelled with DiI (magenta; Allen Brain Atlas: −2.06 mm from Bregma). **c.** Raster plots of an example VPm neuron. This neuron’s spiking decreases to 8 mW/mm^2^, but increases to 30 mW/mm^2^ optogenetic activation of L6CT neurons. **d.** PSTHs of the same VPm neuron in ***c***, indicating that this neuron’s firing rate is bidirectionally modulated by increasing optogenetic activation of L6CT neurons: its firing rate is suppressed relative to baseline to 8 mW/mm^2^, but is enhanced relative to baseline to 30 mW/mm^2^ activation of L6CT neurons. **e.** Grand PSTHs (mean +/− sem across neurons) of the VPm population to 8 and 30 mW/mm^2^ optogenetic activation of L6CT neurons. The population’s ongoing activity is bidirectionally modulated by increasing optogenetic activation of L6CT neurons. **f.** The mean change in ongoing firing rate of the VPm population contributing to the PSTHs in ***e*** is bidirectionally modulated in a graded manner across LED intensities, with a reversal in the trend occurring beyond 8 mW/mm^2^ (+/− sem; 8 mW/mm^2^: −2.06 +/− 1.04 sp/s, p = 1.5e-4; 30 mW/mm^2^: 6.05 +/− 1.68 sp/s, p = 1.3e-3; p-value indicates pairwise comparisons to No LED condition using the two-sided Wilcoxon signed rank test with a Bonferroni correction; ** indicates p < 2e-3; *** indicates p < 2e-4). **g.** Counting the modulatory effect on individual VPm neurons indicates that a majority of neurons are suppressed at 8 mW/mm^2^, whereas a majority of neurons are enhanced at 30 mW/mm^2^ (Change for each neuron was determined by the two-sided Wilcoxon signed rank test on pre- and post-LED input firing rates across trials).

For the example VPm neuron presented in Figures 2c and 2d, 8 mW/mm^2^ optogenetic activation of L6CT neurons results in spiking activity that is persistently below the baseline VPm rate of approximately 6.3 spikes/second. However, at 30 mW/mm^2^, the same VPm neuron’s spiking activity transiently decreases during the ramping portion of the LED input, before increasing and staying above baseline levels for the remainder of the LED input.

Across the population, we found that optogenetic activation of L6CT neurons bidirectionally modulates the ongoing firing rate of VPm neurons. That is, overall, VPm neurons are suppressed relative to baseline at 8 mW/mm^2^ but are enhanced relative to baseline at 30 mW/mm^2^ (Fig 2e). We first characterized the overall effect of the L6CT activation on the VPm by quantifying the population spiking activity during the same “quantification window” used on L6CT neurons, which is during the hold period of the LED input. Across the five presented LED intensities, the change in the ongoing firing rate of VPm neurons is bidirectionally modulated in a graded manner, with relatively low LED intensities suppressing the VPm population firing rate, and a reversal of this suppressive effect occurring at 16 mW/mm^2^, beyond which higher LED intensities enhance the firing rate of the VPm population (Figure 2f).

The data presented here only includes trials with no whisker movement, a factor that is well known to be associated with changes in global brain state and modulation of VPm spiking (Urbain et al., 2015). Analysis of trials with whisker movement shows that the effect of the LED activation of L6CT neurons on the spontaneous activity of VPm neurons is nearly identical for trials with and without whisker movement (Figure S3).

When counting individual neurons, the ongoing firing rate of 60% of recorded neurons (45 of 75) is suppressed at 8 mW//mm^2^, whereas 60% (45 of 75) of the same set of recorded neurons are enhanced at 30 mW/mm^2^ (Figure 2g). This heterogeneity of effects can be observed in simultaneously recorded VPm neurons with preferential responsiveness to the same whisker. Further, bidirectional modulation can be observed in simultaneously recorded neurons with preferential responsiveness to differing whiskers (Figure S4). Thus, likely due to the broad spatial nature of our LED input, the population heterogeneity is not trivially explained by the functional topography of L6CT axonal projections.

Despite L6CT neurons being in a steady-state during the hold period of the LED inputs, spiking in the VPm neuron population is not. This is apparent in Figure 2e where the firing rate in the 8 mW/mm^2^ condition increases slightly during the hold period of the LED input. However, even when restricting the analysis to the last 50 ms of the LED input, the VPm population is still significantly suppressed by the 8 mW/mm^2^ LED input.

### Optogenetic activation of L6CT neurons enhances ongoing firing rates in the TRN

Given that the thalamic reticular nucleus (TRN) is the primary source of L6CT-mediated inhibition in the VPm (Crandall et al., 2015; Golshani et al., 2001; Lam & Sherman, 2010; McCormick & von Krosigk, 1992), it is important to consider the potential role of TRN in our observations. The dynamics of how L6CT neurons enhance TRN spiking, such as the rate of enhancement of TRN spiking relative to the rate of L6CT activation, could play a role in the bidirectional modulation of VPm that results from monotonic activation of the glutamatergic L6CT neurons. To test this, we turned to silicon probe recordings of the spiking activity of populations of individual TRN neurons during optogenetic activation of L6CT neurons (Figure 3a). The identification of TRN neurons recorded at the relevant stereotaxic coordinates was validated by their spike waveform width and measures of their sensory response and modulation from LED activation of L6CT neurons (Figure 3b & S2; Barthó et al., 2014; Ganmor et al., 2010; Guo et al., 2017; Hartings et al., 2003; Lewis et al., 2015; Makinson et al., 2017).

**Figure 3.**
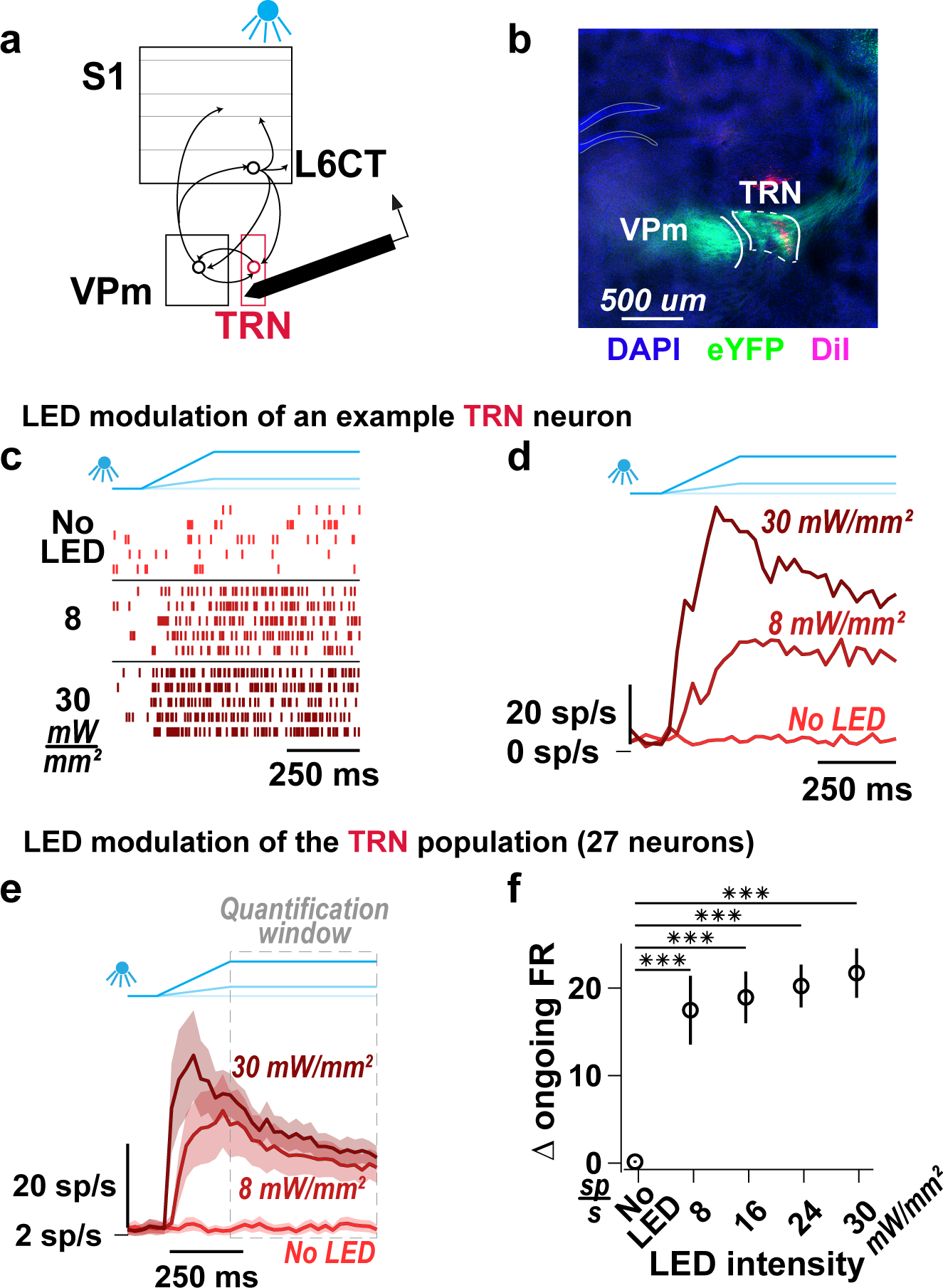
Optogenetic activation of L6CT neurons enhances ongoing firing rates in the TRN. **a.** The direct effect of L6CT neuron activation on TRN is measured from in vivo, silicon probe recordings of neuronal spiking in the somato-sensory section of TRN of awake NTSR1-Cre mice selectively expressing ChR2 in L6CT neurons. **b.** A coronal section of DAPI labelled neurons (blue), ChR2-eYFP expression in L6CT axons terminating in the VPm and TRN (green), and a TRN recording tract labelled with DiI (magenta; Allen Brain Atlas: −1.7 mm from Bregma). **c.** Raster plots of an example TRN neuron. This neuron’s spiking increases to 8 and 30 mW/mm^2^ optogenetic activation of L6CT neurons. **d.** PSTHs of the same TRN neuron in ***c***. **e.** Grand PSTHs (mean +/− sem across neurons) of the TRN population to optogenetic activation of L6CT neurons at 8 and 30 mW/mm^2^. The population’s firing rate is strongly enhanced by both LED intensities. **f.** The mean change in ongoing firing rate of the TRN population contributing to the PSTHs in ***e*** is increased across all LED intensities (+/− sem at 8 mW/mm^2^ = 17.47 +/−3.93 sp/s, p = 5.6e-6; at 30 mW/mm^2^ = 21.71 +/−2.82 sp/s, p = 5.6e-6; p indicates pairwise comparisons to no LED condition using the two-sided Wilcoxon signed rank test with a Bonferroni correction; *** indicates p < 2.5e-4).

The spiking of the example TRN neuron presented in Figures 3c and 3d is strongly increased by the LED at both 8 mW/mm^2^ and 30 mW/mm^2^ (Figure 3e). This strong effect of the LED modulation across all LED intensities largely holds for all recorded TRN neurons (Figure S5). Again, quantifying the population effect during steady state L6CT activation, or the “quantification window”, we observe an enhancement of TRN spiking across all LED intensities. Further, among the LED-on conditions, increasing LED intensity causes relatively small increases in TRN firing rates (Figure 3f). Relative to the monotonic increase in L6CT firing rate with LED intensity that likely induces a similar increase in excitatory conductance in VPm, the inhibitory conductance driven through TRN exhibits a more modest dependence on L6CT activation. This coupled with the strong activation of TRN at the lowest LED intensity likely contributes to the bidirectional influence of L6CT on VPm (see Discussion).

### Optogenetic inputs of increasing LED intensity monotonically enhance the synchrony of the L6CT population

Neurons in circuits with inherently delayed, disynaptic inhibition and highly convergent excitatory synaptic inputs are extremely sensitive to the coordinated timing of those inputs, or the synchrony, making this element of population activity perhaps as important as overall firing rate (Bruno, 2011; Isaacson & Scanziani, 2011; Kremkow et al., 2010; Wang et al., 2010; Wehr & Zador, 2003; Wright et al., 2021). Given the disynaptic inhibition in the L6CT circuit and to further resolve the L6CT-mediated bidirectional changes in VPm ongoing spiking activity, we assessed the synchrony of the L6CT population during our optogenetic manipulations (Figure 4a).

**Figure 4.**
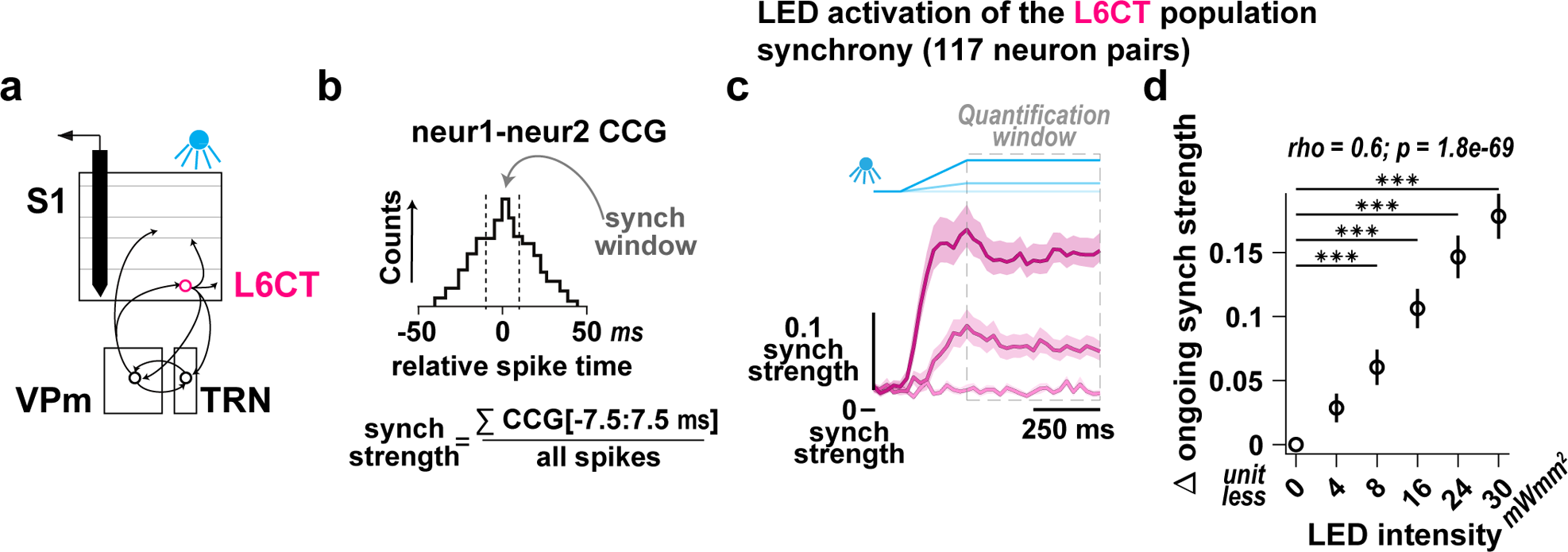
Optogenetic inputs of increasing LED intensity monotonically enhances the synchrony of the L6CT population. **a.** Further assessment of the direct effect of the LED on L6CT neuron activity was measured from the same in vivo, silicon probe recordings of S1 neuronal spiking used in Figure 1. **b.** We assessed the L6CT population synchrony by measuring the strength which is defined for pairs of simultaneously recorded neurons and only includes spikes for a given L6CT neuron that occur within 7.5 ms of another L6CT neuron, normalized by the overall firing rates of the two neurons (see Materials & Methods). **c.** Grand PSTHs (mean +/− sem across neurons) of the L6CT population synchrony strength. The population’s synchrony strength is also enhanced in response to increasing LED intensity. **d.** The mean change in synchrony strength of the L6CT population contributing to the PSTHs in ***c*** is monotonically enhanced by increasing LED intensities (+/− sem; 8 mW/mm^2^: 0.06 +/− 0.01 sp/s, p = 4.6e-5; 30 mW/mm^2^: 0.18 +/− 0.02 sp/s, p = 4.1e-20; p-value indicates pairwise comparisons to no LED condition using the two-sided Wilcoxon signed rank test with a Bonferroni correction; *** indicates p < 2.0e-4; rho & p-value: Spearman’s rank correlation).

Synchronous L6CT spikes are defined as those that occur within 7.5 ms of a spike from another simultaneously recorded L6CT neuron. To quantify synchrony, since firing rate and synchrony covary, we measured the synchrony strength, or the rate of synchronous spikes normalized by the overall firing rate of the pair of neurons (Figure 4b; Liew et al., 2021; Temereanca et al., 2008). The synchrony strength of the L6CT population is also monotonically enhanced by increasing LED intensity. This effect is exhibited in the grand PSTH of the L6CT population synchrony, with a rapid transient increase in synchrony strength during the optogenetic ramp that is sustained during the hold period (Figure 4c). The synchrony strength is summarized over the quantification window in Figure 4d, exhibiting a monotonic relationship with LED intensity. Intriguingly, while we observe a bidirectional trend of the effect of the LED manipulation of L6CT neurons on the ongoing synchrony of the VPm population that parallels the trend of the VPm firing rate presented in Figure 2, the changes are small and nonsignificant (Figure S6). We explore the magnitudes of the changes in L6CT population synchrony strength in subsequent analysis.

### Causal manipulation of L6CT population synchrony results in bidirectional modulation of ongoing firing rates in the VPm

To determine the contribution of L6CT population synchrony to the bidirectional modulation of ongoing VPm activity, for a subset of our recordings (Figure 5a), we designed and presented LED inputs that varied the synchrony of the L6CT population while maintaining a fixed overall firing rate.

**Figure 5.**
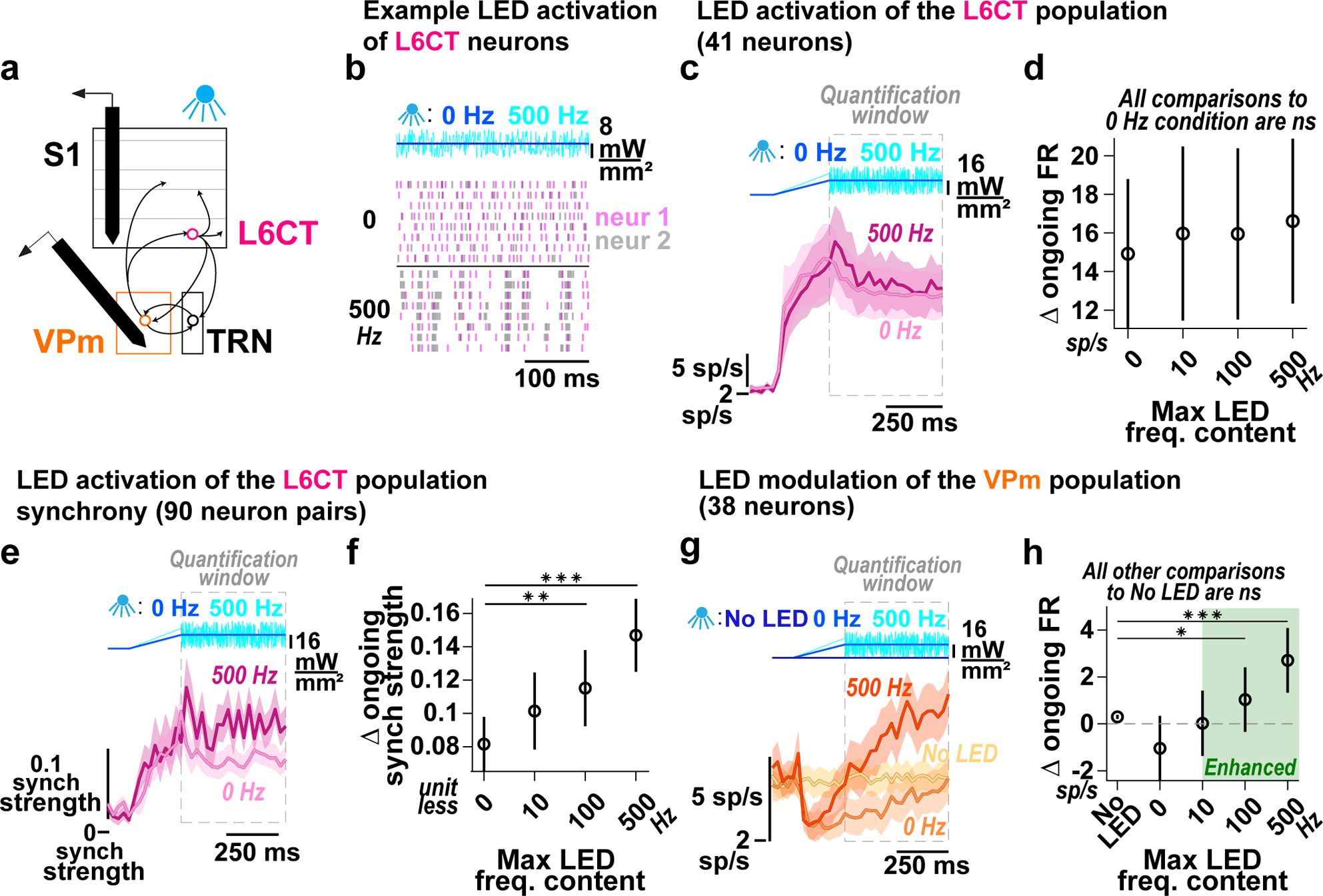
Causal manipulation of L6CT population synchrony results in bidirectional modulation of ongoing firing rates in the VPm. **a.** The effects of additional LED inputs are measured from in vivo, silicon probe recordings of neuronal spiking, either in S1 or the VPm of awake, NTSR1-Cre mice. **b.** Raster plots of two simultaneously recorded L6CT neurons activated by two LED inputs of equal mean, but varying maximum frequency content: 0 or 500 Hz. For better visualization of the time-locked spiking arising from the 500 Hz stimulus, shown are responses to frozen realizations of noise. **c.** Grand PSTHs (mean +/− sem across neurons) of the L6CT population to the 0 and 500 Hz inputs. The L6CT population is similarly activated by the LED inputs. Note that the noise across trials for this analysis is not frozen. **d.** The mean change in ongoing firing rate of the L6CT population contributing to the PSTHs in ***c*** is constant in response to increasing frequency content of the LED input (+/− sem; 0 Hz: 14.92 +/− 3.87 sp/s; 500 Hz: 16.61 +/− 4.27 sp/s, p = 1.2e-1; p-value indicates pairwise comparison to 0 Hz condition using the two-sideed Wilcoxon signed rank test with a Bonferroni correction). **e.** Grand PSTHs (mean +/− sem across neurons) of the L6CT population synchrony. The population’s synchrony strength is enhanced by the 500 Hz stimulus compared to the 0 Hz stimulus. **f.** The mean change in synchrony strength of the L6CT population contributing to the PSTHs in ***e*** is enhanced by increasing frequency content of the LED epoch (+/− sem; 0 Hz: 0.08+/− 0.02 sp/s; 500 Hz: 0.15 +/− 0.02 sp/s, p = 3.8e-9; p-value indicates pairwise comparison to 0 Hz condition using the two-sided Wilcoxon signed rank test with a Bonferroni correction; ** indicates p < 3.33e-3; *** indicates p < 3.33e-4). **g.** Grand PSTHs (mean +/− sem across neurons) of the VPm population to increased frequency content of the LED epoch. **h.** The mean change in ongoing firing rate of the VPm population contributing to the PSTHs in ***g*** is enhanced in a graded manner by the LED frequency content (+/− sem; 0 Hz: −1.18 +/− 1.31 sp/s, p = 1; 500 Hz: 2.97 +/− 1.34 sp/s, p = 3.5e-5; p-value indicates pairwise comparison to No LED condition using the two-sided Wilcoxon signed rank test with a Bonferroni correction; * indicates p < 1.25e-2; *** indicates p < 2.5e-4).

Figure 5b shows the spiking of two simultaneously recorded L6CT neurons in response to two continuous LED inputs: 1) the previously used ramp-and-hold, referred to as the “0 Hz” LED input, or 2) a 250 ms ramp followed by uniformly distributed white noise modulation of the LED intensity with a maximum frequency content of 500 Hz, thus referred to as the “500 Hz” LED input. For this example, frozen realizations of white noise were presented, such that the 500 Hz LED inputs are the same across trials. The 0 and 500 Hz LED inputs have equivalent mean LED intensities and therefore similarly activate L6CT neurons, on average, across trials. However, in response to the 500 Hz LED input, the spike timing across L6CT neurons is often near coincident due to high-frequency events in the LED input. As a result, the rate of synchronous spikes between the pair of neurons is higher compared to that of the 0 Hz LED input.

To further probe the effects of L6CT population synchrony, we presented additional LED inputs consisting of lowpass filtered, non-frozen realizations of white noise with 10 and 100 Hz cutoff frequencies, referred to as the 10 and 100 Hz LED inputs, respectively; all four LED inputs have an equal mean of 16 mW/mm^2^. Across the L6CT population, we find that the firing rate is not significantly changed by the 10, 100, and 500 Hz LED inputs compared to the 0 Hz LED input (Figs. 5c and Figure 5d). However, there is a corresponding, significant increase in the synchrony strength for the noisy LED inputs as compared to the 0 Hz LED input (Figures 5e and 5f). Notably, these synchrony strengths are within the range observed from the ramped steps of light in Figure 4d.

Having now decoupled the L6CT population synchrony from firing rate, we then measured the effect of the increase in L6CT population synchrony on VPm activity. In response to the 0 Hz LED input with mean intensity of 16 mW/mm^2^, the ongoing firing rate in the VPm population is suppressed relative to baseline, although this change is not statistically significant as is also the case in Figure 2f. However, even though the overall firing rate of the L6CT neuron population is the same, increased L6CT synchrony enhances the ongoing firing rate in the VPm population above baseline (Figures 5g and 5h).

Taken together, the synchrony of the L6CT population plays an important role in contributing to the net modulatory effect of L6CT neuron activation on VPm ongoing activity (Figure S7). These shifts in L6CT synchrony can arise from overall changes in L6CT neurons’ firing rates as well as from changes in the temporal precision of equivalent numbers of L6CT neurons’ spikes.

### Modeling the role of excitatory and inhibitory conductances in the L6CT synchrony-dependent bidirectional modulation of VPm ongoing activity

To understand the synaptic conductances underlying the L6CT synchrony-dependent bidirectional modulation of VPm, we constructed a simple circuit model that illustrates the key interactions between L6CT, VPm, and TRN neurons. In contrast to a full network population model matching the numbers of neurons and connections, we followed the spirit of previous reduced complexity models developed for the thalamocortical circuit (Pinto, 2003; Pinto et al., 1996). The circuit consists of two primary nodes capturing the average VPm and TRN population dynamics, each modelled with linear integrate-and-fire (LIF) dynamics. Each node receives common Poisson spike train inputs from 35 L6CT neurons with independently controlled firing rates and synchrony. Each node also receives independent, noisy, excitatory inputs to generate spontaneous spiking. Finally, the TRN node provides inhibitory input to the VPm node (Figure 6a; see Materials and Methods).

**Figure 6.**
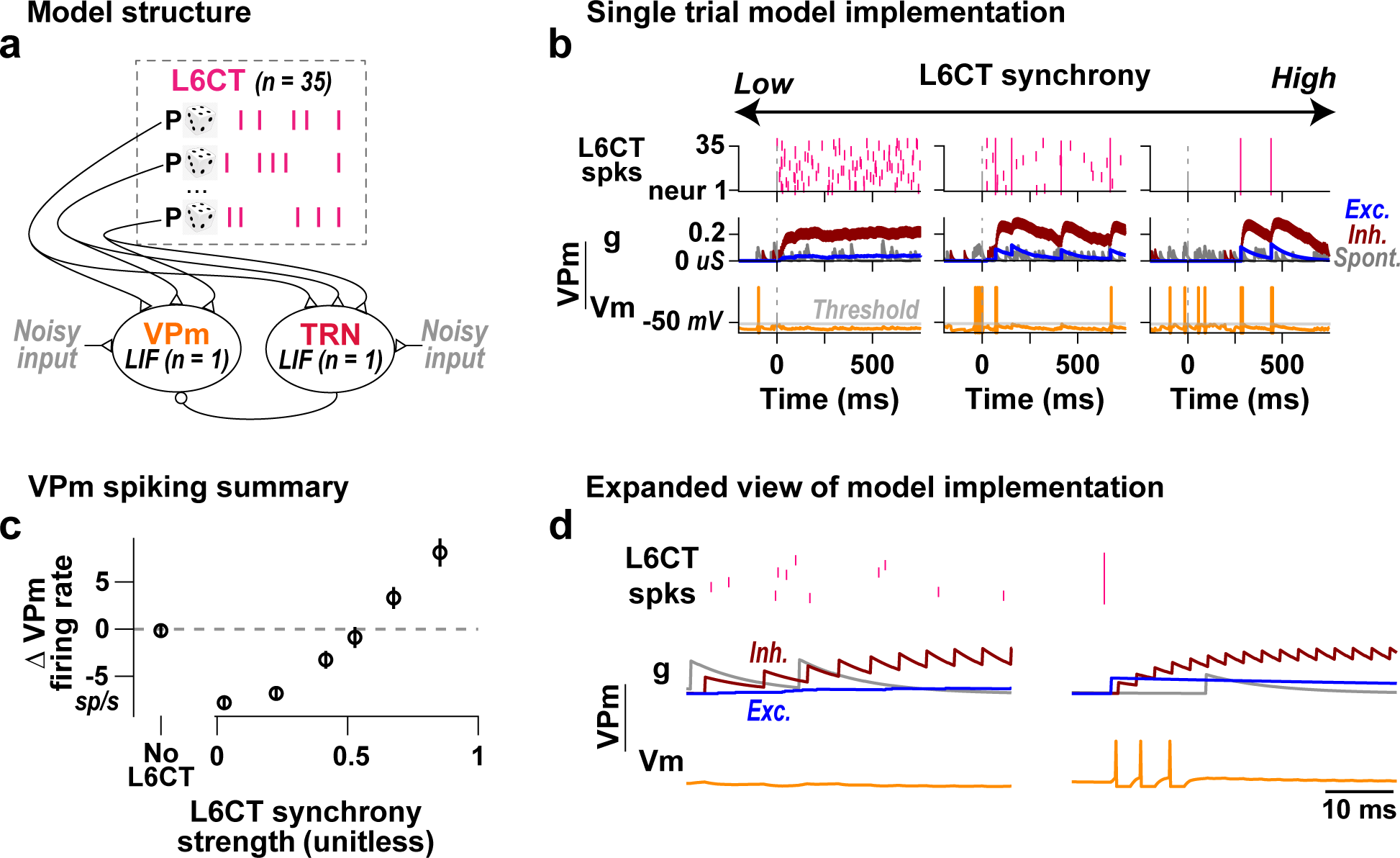
Modeling the role of excitatory and inhibitory conductances in the L6CT synchrony-dependent bidirectional modulation of VPm ongoing activity. **a.** Representative VPm and TRN neurons were modeled as leaky, integrate and fire (LIF) neurons, each integrating common L6CT spike train inputs with controlled synchrony and firing rate. The VPm neuron integrated inhibitory input from the spiking of the TRN neuron. Spontaneous activity for each LIF neuron was driven by noisy, excitatory inputs. **b.** Single trial model simulations run by holding the L6CT neurons’ firing rates fixed while varying the synchrony of the L6CT input result in a shift from suppression to enhancement of VPm spiking in a graded manner with increasing synchrony. Note that the spontaneous conductance has been scaled 5x for better visualization. **c.** The mean (+/− sem across 50 trials) change in VPm firing rate with increasing L6CT population synchrony, but constant L6CT neurons’ firing rate (2.5 sp/s) replicates the experimentally observed, L6CT synchrony-dependent, bidirectional modulation of VPm. **d.** An expanded view of the single trial components and the underlying conductances in VPm highlights the key factors underlying L6CT synchrony-dependent, bidirectional modulation of VPm: (1) the relative strength of the inhibitory conductance compared to the excitatory conductance, and (2) the relative timing of the monosynaptic excitatory and disynaptic inhibitory conductances.

Figure 6b shows single trial realizations of the model, where we assess the conductances and membrane potential in the VPm as we increase the synchrony of the L6CT spike trains while maintaining a constant firing rate. Here, the excitatory conductance arises from L6CT spiking and the inhibitory conductance arises from TRN spiking.

VPm spiking activity replicates our experimental results: a suppression of spontaneous spiking at low synchrony and an enhancement of spiking at high synchrony. Quantifying the average change in VPm firing rate across trials, we observe an L6CT synchrony-dependent bidirectional modulation of VPm activity across a continuum of L6CT synchrony strengths (Figure 6c). Unsurprisingly, the model produces much less variability across trials than the experimental data. Notably, these synchrony strengths are within the range observed in Figure 4d and reiterate that even seemingly small changes in measures of synchrony can have notable effects on the firing activity of downstream neurons (Wright et al., 2021).

To understand the evolution of the membrane potential under varying levels of L6CT synchrony, we examined the underlying model conductances, an aspect that is hidden in our extracellular recordings. We found that under the low synchrony condition, the magnitude of the inhibitory conductance dominates over that of the excitatory conductance for the entire trial duration. Individual L6CT spikes cannot sufficiently activate the excitatory conductance to ultimately depolarize the membrane potential for spiking. Moreover, not only do individual L6CT spikes sufficiently activate the inhibitory conductance, but the inhibitory conductance never decays back to baseline due to the frequent occurrence of L6CT spikes. As a result, the VPm membrane potential is hyperpolarized and spontaneous spiking is suppressed (Figure 6d). In contrast, in the high synchrony condition, the synchronous L6CT spikes result in an excitatory conductance that is sufficient to enable summation to spiking before the onset of the disynaptic, inhibitory conductance.

There are several well-documented factors that undoubtedly play a role in our experimental findings, such as the short-term dynamics of the involved synapses (Crandall et al., 2015) as well as the fact that L6CT inputs result in modulatory NMDA plateau potentials in the distal dendrites of FO thalamic neurons (Sherman & Guillery, 1998). We propose that the interplay of both the relative strength and timing of the excitatory and inhibitory conductances, as observed in our model, is a relatively simple mechanism that plays an additional key role in our experimental findings. In this scenario, the synchrony of the L6CT neuron population strongly influences the net modulatory effect of L6CT neuron input on its VPm neuron target, thus serving as an important factor in L6CT function.

### Optogenetic activation of L6CT neurons enhances VPm sensory responses

Having established that different levels of activation of L6CT neurons cause bidirectional changes in the ongoing firing rate in VPm neurons, we next sought to understand the effect of L6CT neuron activation on sensory responses evoked in the VPm. We recorded the spiking response of populations of individual VPm neurons to concurrent single whisker deflection and optogenetic L6CT input of increasing LED intensity (Figure 7a).

**Figure 7.**
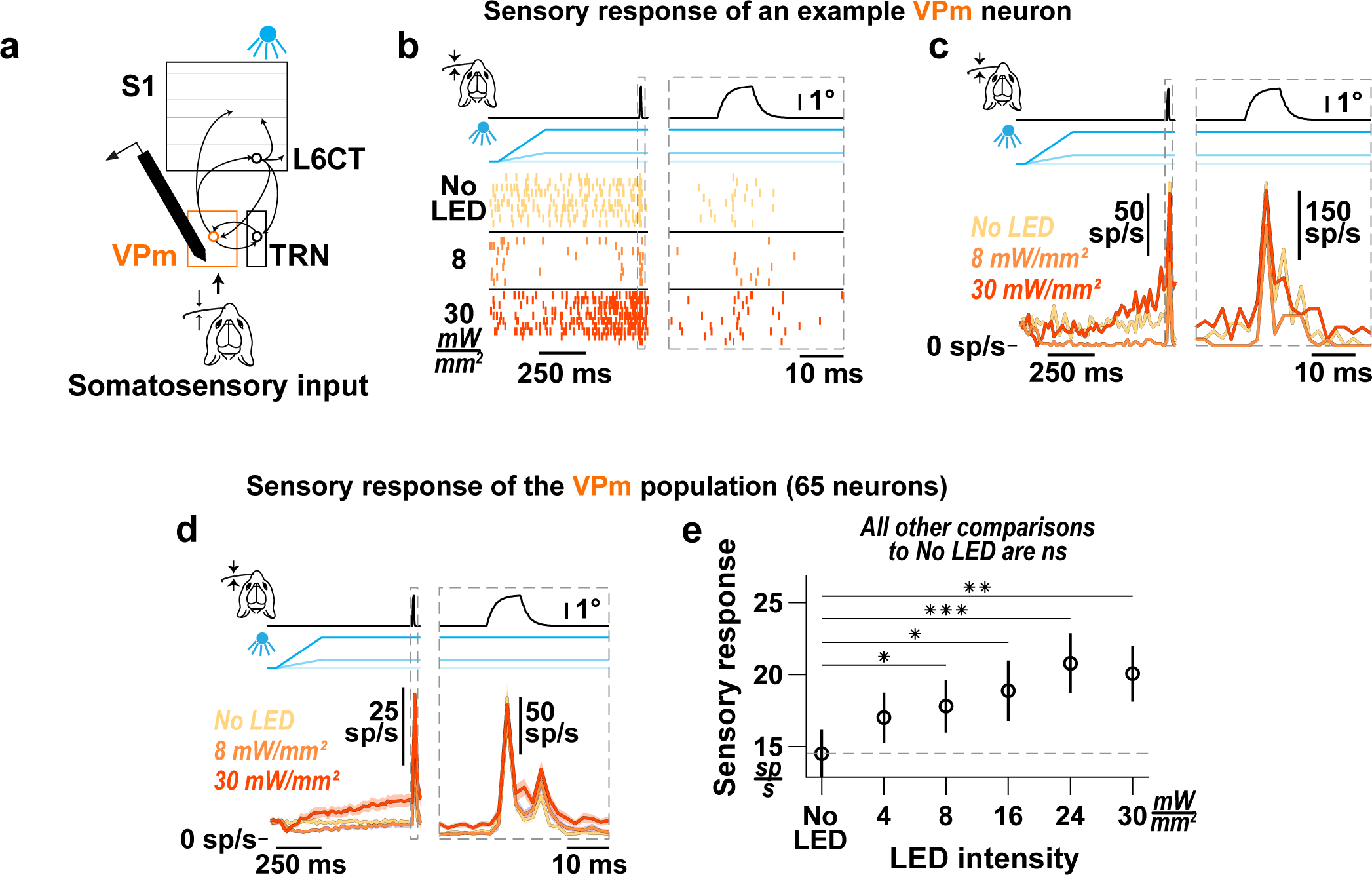
Optogenetic activation of L6CT neurons enhances VPm sensory responses. **a.** The effect of L6CT neuronal activation on sensory responses in VPm is measured from recordings of neuronal spiking in the VPm while concurrently deflecting a single whisker for somatosensory input and increasing optogenetic activation of L6CT neurons in awake, NTSR1-Cre mice. **b.** Raster plots of an example VPm neuron. This neuron’s ongoing spiking activity is bidirectionally modulated and its stimulus-evoked response to sawtooth, somatosensory input is enhanced by increasing optogenetic activation of L6CT neurons. The right panel is an expanded view of the same neuron’s sensory response. **c.** PSTHs of the same neuron’s spiking in ***b***. Note the change in the vertical scale in the right and left views due to differing bin sizes. **d.** Grand PSTHs (mean +/− sem across neurons) of the VPm neuron population. The population’s ongoing activity is bidirectionally modulated, but the stimulus-evoked response is enhanced by increasing optogenetic activation of L6CT neurons. The right panel is an expanded view of the same population’s sensory response. Note the change in the vertical scale in the right and left views. **e.** Optogenetic activation of L6CT neurons increases the mean, baseline-subtracted, stimulus-evoked response of the VPm population contributing to the PSTHs in ***d*** (+/− sem; No LED: 14.47 +/− 1.59 sp/s; 8 mW/mm^2^: 17.7 +/− 1.76 sp/s, p = 8.2e-3; 30 mW/mm^2^: 19.95 +/− 1.87 sp/s, p = 5.3e-4; p-value indicates pairwise comparison to No LED condition using the two-sided Wilcoxon signed rank test with a Bonferroni correction; * indicates p < 1.0e-2; ** indicates p < 2.0e-3; *** indicates p < 2.0e-4).

Figures 7b and 7c show the response of an example VPm neuron to a sawtooth whisker stimulus occurring 750 ms after the onset of the LED activation of L6CT neurons. As previously noted, the effect of the optogenetic activation of L6CT neurons on ongoing activity in VPm neurons varies with LED intensity, with a suppression at 8 mW/mm^2^ and an enhancement at 30 mW/mm^2^ (Figure 7b, left panels). Taking the direction of change in ongoing firing rate as a cue for the general excitability of VPm neurons, a simple prediction would be that the sensory response in VPm neurons would be amplified when VPm background activity is increased and attenuated when VPm background activity is suppressed. However, as qualitatively highlighted by the example VPm neuron sensory response in Figures 7b and 7c, despite bidirectional modulation of the ongoing activity of this neuron, the sensory response is not bidirectionally modulated by the LED inputs.

We accounted for both the pre- and post-sensory stimulus modulatory effects of L6CT activation and quantified the sensory response by measuring the trial-averaged stimulus-evoked rate subtracted from the pre-stimulus rate, in equivalent 30 ms duration windows. Across the entire VPm population, increasing optogenetic activation of L6CT neurons tends to enhance the magnitude of the sensory response (Figures 7d and 7e). These effects are measured in trials without whisker movement, as presented in Figure 7. The effect in trials with whisker movement is associated with a reduction of the overall magnitude of the VPm sensory response as well as a reduction in the measured effect of L6CT activation on the VPm sensory response (Figure S8). We also observe a corresponding increase in the sensory response synchrony strength of the VPm population response across all LED intensities (Figure S9).

Interestingly, the VPm population sensory response is not further affected when we present the sensory stimulus during the 500 Hz noise LED input, as compared to the 0 Hz LED step of equal mean. This suggests that a selective increase in the synchrony of the L6CT inputs does not further impact the sensory responses (Figure S10), in contrast to what is observed for ongoing VPm activity (Figure 5).

To assess the effect of L6CT modulation on the transfer of sensory inputs through VPm in a slice preparation, Crandall et al. co-activated L6CT neurons and medial lemniscus (ML) axons. They found a clear relationship between the effect of L6CT activation on the response of VPm neurons to ML stimulation and the biphasic modulatory effect on spontaneous VPm spiking (Crandall et al., 2015). From such a study, it is reasonable to expect that VPm sensory responses would be suppressed in conditions where VPm ongoing activity is suppressed, at 4 and 8 mW/mm^2^. Yet, we rather observe that neurons with increasingly suppressed ongoing activity tend towards a more enhanced sensory response compared to the no LED condition (Figure S11). While one might expect an increase in T-type calcium channel-mediated burst spikes in conditions where the VPm ongoing activity is suppressed (Borden et al., 2022; Kirchgessner et al., 2020), we do not observe any significant changes in stimulus-evoked VPm bursting upon L6CT activations (Figure S12). Thus, the observed bidirectional changes in VPm ongoing activity do not trivially predict the VPm sensory response magnitudes, as sensory responses are consistently amplified by L6CT activation.

### Optogenetic activation of L6CT neurons dynamically modulates activity in S1

Given the effects of optogenetic activation on ongoing and stimulus-evoked activity in VPm neurons, it is important to determine how these cortico-thalamo-cortical modulatory effects coupled with modulatory effects directly from L6CT intracortical projections ultimately impact S1. We recorded the spiking response of populations of individual S1 neurons across all cortical layers while presenting the same optogenetic and tactile inputs (Figure 8a). All neurons were identified as either putative excitatory regular-spiking (RS) or inhibitory fast-spiking (FS) neurons based on the trough to peak time of their spike waveform widths (Figure 8b; See Materials and Methods).

**Figure 8.**
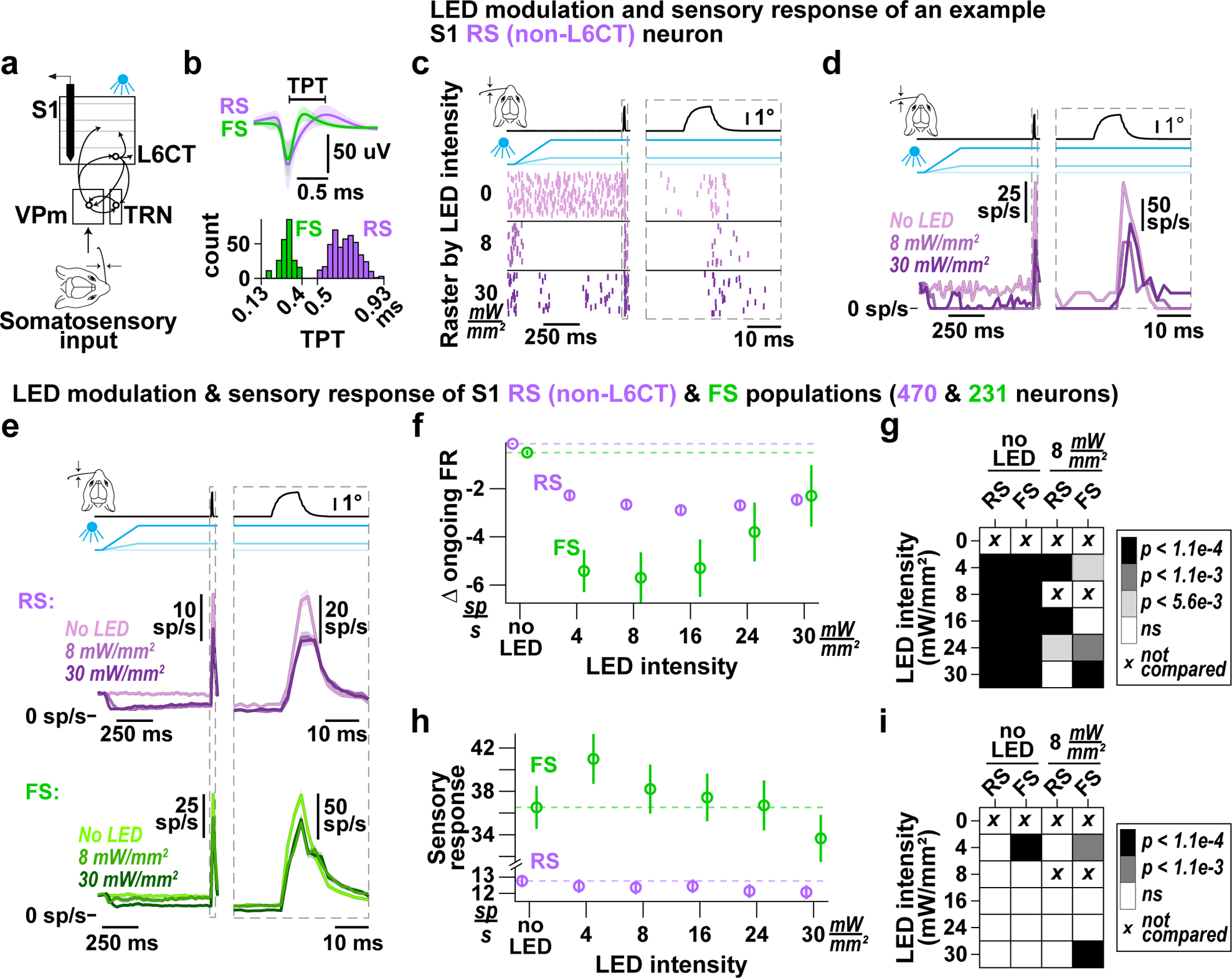
Optogenetic activation of L6CT neurons dynamically modulates activity in S1. **a.** The effect of L6CT neuron activation on ongoing activity and sensory responses in S1 is measured from recordings of neuronal spiking in S1 while concurrently deflecting a single whisker for somatosensory input and increasing optogenetic activation of L6CT neurons in awake, NTSR1-Cre mice. **b.** Mean waveform +/− standard deviation across S1 neurons identified as either regular spiking (RS) or fast spiking (FS) based on the trough to peak time (TPT) of their spike waveform (RS > 0.5 ms and FS < 0.4 ms; see Materials and Methods). **c.** Raster plots of an example RS (non-L6CT) neuron. This neuron’s ongoing and stimulus-evoked spiking is suppressed by concurrent optogenetic activation of L6CT neurons (left panel). The right view is an expansion of the same neuron’s sensory response. **d.** PSTHs of the same neuron’s spiking in ***c***. Note the change in the vertical scale in the right and left views due to differing bin sizes. **e.** Grand PSTHs (mean +/− sem across neurons) of the S1 RS (non-L6CT) and FS neuron sub-populations from all cortical layers. The right view is an expansion of the population sensory response; note the change in the vertical scale in the right and left views. **f.** The mean +/− sem change in ongoing activity of the S1 RS (non-L6CT) and FS sub-populations contributing to the PSTHs in ***e***. Both sub-populations are suppressed by all LED inputs, although to different extents (RS: 8 mW/mm^2^: −2.67 +/− 0.2 sp/s; 30 mW/mm^2^: −2.49 +/− 0.25 sp/s. FS: 8 mW/mm^2^: −5.68 +/− 1.06 sp/s; 30 mW/mm^2^: −2.26 +/− 1.28 sp/s). Additionally, there is an apparent reversal, in the suppressive effect beyond 16 mW/mm^2^, reminiscent of the reversal effects observed in VPm ongoing activity. **g.** Confusion matrix indicating statistical significance in the change in ongoing firing rate observed in ***f*** for each sub-population when comparing to both no LED conditions as well as to 8 mW/mm^2^ (first and last two columns, respectively); color indicates p-value using the two-sided Wilcoxon signed rank test with a Bonferroni correction. Both sub-populations are significantly suppressed across LED intensities compared to no LED levels. In addition, the change at 8 mW/mm^2^ is significantly lower than at 4, 16, and 24 mW/mm^2^ for the RS sub-population and 4, 24, and 30 mW/mm^2^ for the FS sub-population. **h.** Same as ***f***, but for sensory responses. While the RS sub-population is modestly suppressed across all LED intensities, there is an increase of the FS sub-population sensory response at 4 mW/mm^2^ (RS: No LED: 13.15 +/− 0.58 sp/s; 8 mW/mm^2^: 12.55 +/− 0.62 sp/s; 30 mW/mm^2^: 12.16 +/− 0.64 sp/s. FS: No LED: 36.62 +/− 1.98 sp/s; 4 mW/mm^2^: 41.1 +/− 2.3 sp/s; 8 mW/mm^2^: 38.33 +/− 2.26 sp/s; 30 mW/mm^2^: 33.78 +/− 2.16 sp/s). However, this enhancive effect is reversed at higher intensities. **i.** Same as ***g***, but for sensory responses. While there is no significant change in sensory response in the RS sub-population, the sensory response for the FS sub-population is significantly enhanced at 4 mW/mm^2^ compared to no LED conditions. Further, compared to the response at 8 mW/mm^2^, the response at 4 mW/mm^2^ is significantly enhanced and the response at 30 mW/mm^2^ is significantly suppressed.

Figures 8c and 8d show an example S1 RS (non-L6CT) neuron’s response to optogenetic activation of L6CT neurons at varying LED intensities with an embedded sensory stimulus. Pre-stimulus, the LED clearly suppressed the ongoing spiking activity of this neuron. Intriguingly, more of the spiking was suppressed at 8 mW/mm^2^ than at 30 mW/mm^2^. As shown in the right panels of Figures 8c and 8d, the sensory response of this neuron was also modestly suppressed by optogenetic activation of L6CT neurons.

Both when looking at RS and FS sub-populations across all layers (Figs. 8e & 8f), as well as when parsing neuronal responses by layers (Figure S13 & S14), optogenetic activation of L6CT neurons leads to a suppression of ongoing activity in both S1 sub-populations, with FS neurons suppressed to a larger extent. This suppression likely arises in large part from L6CT direct intracortical projections to S1 inhibitory neurons, as previously characterized in the visual cortex (Bortone et al., 2014).

While there is a clear suppression of both sub-populations at 8 mW/mm^2^, there is no further suppression at higher LED intensities. Rather, in the RS sub-population, there is a small reversal of the suppressive effects beyond 16 mW/mm^2^, reminiscent of the observed changes in ongoing VPm activity that would then be propagated to S1 (Figure 8f). To assess this reversal, we made pairwise comparisons between the 8 mW/mm^2^ condition and all other LED-on conditions. While the reversal is small in the RS sub-population, the change in ongoing firing rate in the FS population is significantly higher at 24 and 30 mW/mm^2^ compared to the change at 8 mW/mm^2^ (Figure 8g). Such significance is also observed in FS populations in layers 4 and 5, such that the change in ongoing activity in response to the 30 mW/mm^2^ condition is statistically unchanged from baseline compared to the clear suppression observed in response to the 4 and 8 mW/mm^2^ inputs (Figure S13b). Despite the well-documented effects of behavioral state changes reflected in whisker movement on overall cortical spiking (reviewed in Poulet & Crochet, 2019), whisker movement was associated with only a modest decrease in the measured effect of L6CT activation on ongoing RS activity and was not associated with any change in the effect of L6CT activation on ongoing FS activity (Figure S15).

We then characterized the effect of optogenetic activation of L6CT neurons on sensory responses of the S1 sub-populations. While there is an overall small decrease in the RS (non-L6CT) population sensory response between the no LED condition to LED-on conditions (less than 1 sp/s; Figure 8h), the sensory response for most S1 RS (non-L6CT) neurons is statistically unchanged, such that all pairwise comparisons of the RS population’s sensory response between no LED conditions to LED-on conditions of increasing intensities are nonsignificant (Figure 8i). Whisker movement is associated with an overall decrease in RS sensory responses across all LED intensities. Further, pairwise comparisons of the RS population’s sensory response between no LED and LED-on conditions are also nonsignificant in trials with whisker movement (Figure S16). Moreover, when parsing RS sensory responses by layer, the only clear trend that emerges is in layer 6, where responses are suppressed (Figure S17b).

Interestingly, the sensory response of FS neurons at 4 mW/mm^2^ is significantly higher than baseline conditions, whereas the response at 30 mW/mm^2^ is non-significantly suppressed. Given the observed maximal suppression of ongoing FS activity at 8 mW/mm^2^ as well as the apparent reversal in the effect of LED intensity on sensory response, we again compared the sensory response of all LED-on conditions to that at 8 mW/mm^2^. Under this analysis, the sensory response at 30 mW/mm^2^ is significantly lower than the response at 8 mW/mm^2^ (Figure 8i). Features of these findings are present when parsing the neuronal response by layer, with significance in layers 4 and 5 (Figure S18). Despite an overall decrease in sensory response across all LED intensities in trials with whisker movement, the trend of an enhanced response at 4 mW/mm^2^ and suppressed response at higher intensities is also present but nonsignificant in trials with whisker movement (Figure S16).

The net observed effect of corticothalamic feedback on S1 is a sum of both L6CT-mediated changes through the cortico-thalamo-cortical route of modulation as well as through direct intracortical projections. This complexity underlies our results. While the overall effect of optogenetic activation of L6CT neurons on ongoing activity in both S1 subpopulations is suppressive, the effect on the sensory responses of the FS sub-population is dynamic, such that there is a sweet spot of enhancement. These dynamics would then contribute to the observed response in the RS population.

## DISCUSSION

In this study, we conducted a functional dissection of the somatosensory cortico-thalamo-cortical circuit by way of multi-region electrophysiology, strategic design of optogenetic manipulations and sensory stimulation, as well as a simple circuit model. In doing so, we determined the direct effect of L6CT neurons on thalamocortical sensory processing across a range of activation patterns during wakefulness. We expanded on previous in-vivo findings that provide evidence for L6CT neurons as static modulators that either suppress *or* enhance FO thalamus (Born et al., 2021; Denman & Contreras, 2015; Guo et al., 2017; Mease et al., 2014; Olsen et al., 2012; Pauzin & Krieger, 2018; Spacek et al., 2022; Temereanca & Simons, 2004; Voigts et al., 2020) and establish L6CT feedback as a dynamic modulator that can both suppress *and* enhance the ongoing spiking of a VPm neuron. Further, we pinpoint that the bidirectional modulation of VPm is mediated both by differential sensitivity of L6CT and TRN neurons to increasing LED inputs as well as specific aspects of the activity patterns of the L6CT population: the firing rate and synchrony, properties that, generally, vary in the cortex across behaviors (Poulet & Crochet, 2019). In addition, we provide evidence for the effect of this dynamic modulation on S1 neurons and find non-monotonic changes in S1 RS and FS ongoing activity as well as FS stimulus-evoked activity. Collectively, L6CT neurons play a role in the continuous and context-dependent modulation of thalamocortical sensory signaling through motor corollary signals, arousal, and other behavioral states (Augustinaite & Kuhn, 2020; Clayton et al., 2021; Dash et al., 2022; Reinhold et al., 2023; Vélez-Fort et al., 2018), and we provide comprehensive evidence for the effect of the L6CT feedback mechanism across such facets.

Much of what we know about this complex circuit has been learned from studies conducted in ex-vivo brain slice preparations examining L6CT feedback at the cellular level (Crandall et al., 2015; Liu et al., 1995; McCormick & von Krosigk, 1992, 1992; Sherman & Guillery, 1998, 2002; Sherman & Koch, 1986b; Usrey & Sherman, 2019; Van Horn et al., 2000). The consensus from these studies is that synaptic input from L6CT projections serves to modulate the excitability of FO thalamic neurons. But conflicting results have pointed to both excitatory and suppressive effects on excitability. Importantly, in an ex-vivo slice preparation, Crandall et al. found that L6CT neurons can switch their influence on each VPm neuron from excitatory to suppressive depending on the frequency of stimulation (Crandall et al., 2015). The general nature of those results is supported here, but the slice experiments pointed to modulation that is mediated by short-term synaptic plasticity. The L6CT synchrony effects demonstrated here likely work synergistically with the synaptic plasticity mechanisms at the thalamo-reticular targets to provide a wide and continuous range of activity-dependent, top-down modulation of VPm.

The strong activation of TRN at low intensities coupled with the rate of increased activation of L6CT spiking with increased LED intensity compared to the rate of increased activation of TRN spiking likely also plays a role in the bidirectional modulation of VPm. More specifically, the strong activation of TRN at even relatively low levels of optogenetic activation of L6CT (4 mW/mm^2^) could explain an overall suppression of VPm spiking. But the effect of increasing light intensity on L6CT versus TRN spiking suggests the excitatory conductance to VPm is likely increasing at a relatively faster rate than the inhibitory conductance with increased LED intensity, the net sum of which would contribute to the observed bidirectional modulation of VPm.

As suggested by the circuit model in this study, the L6CT synchrony-dependent, bidirectional modulation of VPm emerges, first, because of the strong inhibitory conductance from TRN inputs relative to the strength of the direct excitation from L6CT neurons. The second important feature is the circuit architecture that enables a monosynaptic excitatory conductance and an inherently delayed, disynaptic inhibitory conductance in the VPm. These elements are key to the “window of opportunity” classically described in the feedforward path for FO thalamic inputs onto layer 4 cortical RS neurons (Bruno, 2011; Isaacson & Scanziani, 2011; Pinto, 2003). The premise of this mechanism is that while the strength of individual thalamocortical synaptic inputs is modest, numerous inputs converge on cortical synaptic targets, making the cortical neurons highly sensitive to the synchrony of the ascending inputs (Bruno, 2011). Indeed, L6CT inputs to VPm and TRN are numerous and highly convergent (Guillery & Sherman, 2002; Liu et al., 1995; Sherman & Koch, 1986b; Van Horn et al., 2000), contributing to a synchrony-dependent effect of this classically modulatory circuit. This would provide another example to support the possibility of a synchrony-driven window of opportunity as a canonical circuit computation in the brain.

It is, however, important to note that there are several fundamentally different biological considerations between L6CT and thalamocortical inputs that would impact the window of opportunity in the feedback versus feed-forward circuits. First, L6CT neurons have a wide range of axon conduction latencies relative to that of thalamocortical projections. This would impact the synchronization of the post-synaptic conductances towards summation in the sub-cortical targets. It is worth noting that latencies for corticothalamic signal propagation in the range of 1-7 ms have been measured amongst the full range of axon conduction latencies (reviewed in Briggs & Usrey, 2008). This would lead to a variability that is relatively small compared to the 15 ms (+/− 7.5 ms) window of opportunity that we consider here. Nonetheless, the full impact of the heterogeneity of L6CT conduction velocities in our study is currently unknown and should be pursued in future studies. In addition, it is well known that thalamocortical inputs produce fast conductance changes on the proximal dendrites of their postsynaptic targets via ionotropic receptors. However, L6CT neurons activate both ionotropic and metabotropic receptors, the latter of which produces prolonged NMDA plateau potentials in the distal dendrites of FO thalamic neurons (Sherman & Guillery, 1998). While the time constants of the L6CT synapses in our model were adjusted to reflect the longer timescale of NMDA plateau potentials (Table 1), the nature of our single-compartment model precludes capturing such biological complexity. It is nonetheless important to note that such factors undoubtedly contribute to our experimental results and would impact the window of opportunity. Also, both the L6CT synapses onto VPm and TRN as well as the TRN synapses onto VPm undergo short-term synaptic plasticity (Crandall et al., 2015). None of these factors were implemented in the circuit model in this study because our goal was to construct the simplest possible model to explain the observed phenomena. Undoubtedly though, the dynamics of these synapses are changing in the experimental conditions utilized here and likely complement the demonstrated synchrony-dependent effects. Finally, VPm and TRN neurons have connectivity between each other, in addition to recurrent connectivity within TRN. While the initial transient response to the optogenetic activation engages the full range of connectivity in this circuit, the later steady-state response appears to be dominated by L6CT-VPm and L6CT-TRN-VPm projections. In the model, the TRN-VPm influence likely reflects the combined effects of the omitted circuit elements that together set the overall inhibitory tone and dictate the window of opportunity.

The very nature of the reciprocal connectivity of the thalamus and cortex makes assigning causal influences challenging. While optogenetic interventional approaches in the primary visual cortex have shown that the suppression of the ongoing activity of both the RS and FS S1 subpopulations arises predominantly from direct intracortical L6CT projections to inhibitory translaminar cells (Bortone et al., 2014), the cortical sensory response reflects the net culmination of two sub-circuits: the intracortical circuit and the cortico-thalamo-cortical circuit. The net effect of the L6CT activation on ongoing, baseline activity of both RS and FS neurons across cortical S1 is suppressive. The larger suppression of the cortical FS neurons relative to RS points to a change in E/I balance and a corresponding increase in cortical excitability with L6CT activation. Although the baseline-subtracted sensory-evoked response of the cortical RS neurons showed no apparent change with L6CT activation, the suppression of ongoing activity could result in a more potent effect of this population activity on downstream targets through a decrease in levels of synaptic depression and/or changes in the signal-to-noise ratio of the sensory stimulus.

Of the most dramatic effects shown here is the effect of the L6CT activation on S1 FS neurons, which exhibits a “sweet spot” of an enhanced cortical sensory-evoked response, which would then additionally inhibit cortical RS neurons. The result of all these sub-circuits being modulated by L6CT neurons is an RS sensory response magnitude that is surprisingly invariant to the manipulation. However, the effects of the differential modulation of the RS and FS sub-populations could have a profound impact on spatiotemporal sensory representations. Additional experimental investigation would be required to fully disentangle the contributions of each cortico-thalamo-cortical and intracortical sub-circuits, such that contributions from each individual component could be selectively and reversibly toggled. Moreover, while overall firing rates in RS neurons were unchanged, it is possible that L6CT activation could dynamically modulate other, more context-dependent aspects of RS neurons, such as their encoding of deviant (Voigts et al., 2020) or adapting (Mease et al., 2014) stimuli, as has been explored in recent studies.

The experimental design likely has some effects on the observations made in this study. First, the sensory stimulus consisted of a brief, punctate, whisker deflection. The impulse-like nature of the stimulus is advantageous for establishing the input-output properties within a circuit. Moreover, the punctate stimulus is akin to stick-slip events that occur when a mouse palpates a surface, and thus is a relevant stimulus for probing sensory processing in the whisker pathway (Jadhav et al., 2009; Wolfe et al., 2008). However, it is entirely possible that the effect of L6CT neuron activation on a more sustained and persistent sensory stimulus would be quite different. In contrast to the punctate sensory inputs utilized here that drive synchronous VPm spiking which would then strongly drive S1, sequences of whisker deflections such as those occurring during texture or curvature discrimination (Jadhav et al., 2009; Rodgers et al., 2021) or continuous noisy whisker inputs could result in more measurable modulation of the sensory response of cortical RS neurons than observed here.

Second, the optogenetic activation of the L6CT neurons was achieved through LED illumination of the cortical surface in a broad and diffuse manner. While L6CT neuron projections are topographically organized, TRN projections are not, which results in modulatory effects in the FO thalamus that depend on topographic alignment (Born et al., 2021; Temereanca & Simons, 2004). However, the spiking activity of L6CT neurons has been shown to be modulated by extra-sensory features (Augustinaite & Kuhn, 2020; Clayton et al., 2021; Vélez-Fort et al., 2018). In those cases, the activation of L6CT neurons would likely be global, or at least extend to multiple barrel columns within S1, which is the type of conditions that the optogenetic manipulation here would represent. Moreover, the increase in L6CT neuron spiking would likely be long-lasting, more akin to sustained, long-duration manipulation of activation here, as opposed to optogenetic pulsing or analysis of epochs after the optogenetic activation of L6CT neurons utilized in other studies (Guo et al., 2017; Kirchgessner et al., 2020). Such experiment designs likely modulate the circuit in an entirely different manner and explain differences in our study and others, such as a lack of changes in FO thalamic bursting observed here.

Finally, our experiments were conducted in head-fixed, stationary mice, such that the natural activity range of L6CT neurons would preclude that arising from many extrasensory behavioral states. Our study lays the foundation of the direct effect of L6CT feedback on thalamocortical activity across a range of L6CT activity. Future studies must continue to characterize L6CT neurons (both their firing rates and synchrony) across a variety of behaviors and then manipulate L6CT neurons to causally assess the contribution of L6CT neuron spiking to these behaviors. Although architectures involving loops provide challenges to assigning causality, parsing observations based on behavioral/brain states and/or causal manipulation through interactive, closed-loop adaptive experimental strategies (Bolus et al., 2018, 2021) may provide a path to disentangle complex circuits such as these involved in sensorimotor behaviors.

In our world that is full of dynamically changing sensory inputs, it is crucial for survival to be able to flexibly adjust our responsiveness to these inputs. Under contexts different from that studied here, Guo et al. found that the dynamic regulation facilitated by L6CT neurons in FO thalamus mediates a trade-off between the detectability and discriminability of auditory stimuli (Guo et al., 2017). Indeed, the dynamic effect of L6CT neurons and their location relatively early in the sensory processing hierarchy positions L6CT neurons to play critical roles in sensory processing, bidirectionally modulating activity in a way that would serve a variety of behavioral needs. Further, in addition to sensory circuits, the L6CT neuron circuit motif is also present in motor systems (Galvan et al., 2016), potentially pointing to a canonical circuit motif that subserves dynamic neural signaling in complex environments.

## Acknowledgements

We thank Simone Russo and Kayla Peelman for their helpful comments on the manuscript. We thank Barry Connors’ laboratory at Brown University for providing us with the NTSR1-cre mouse line.

---

**Figure S1.**
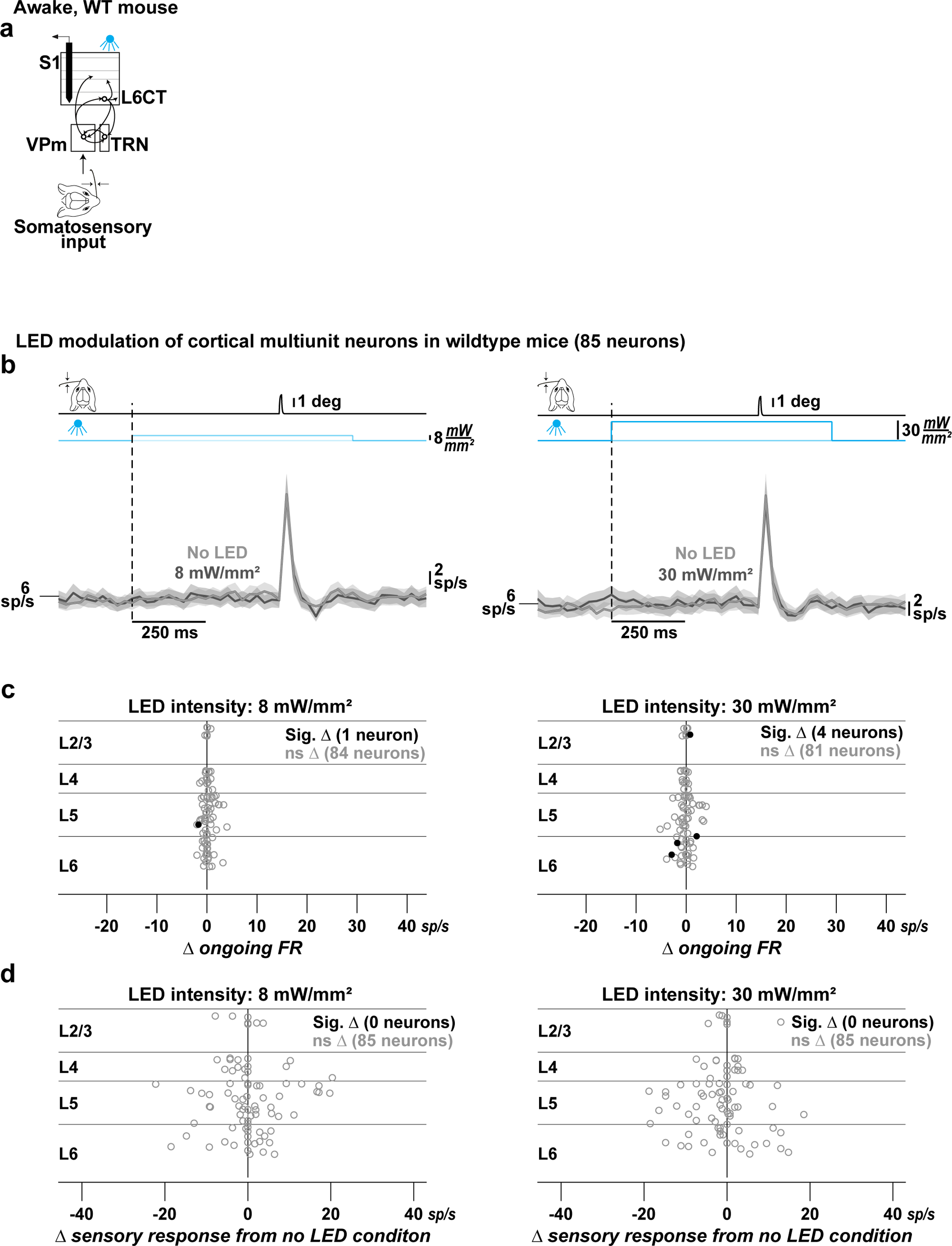
The LED input alone does not explain our observed effects of L6CT activation on the cortico-thalamocortical circuit. **a.** The direct effect of the LED input alone on ongoing and stimulus-evoked cortical neuron activity is measured from in vivo, silicon probe recordings of neuronal spiking in S1 of an awake, head-fixed, wildtype (WT) mouse with no opsin expression. As in the experiments with opsin-expressing, NTSR1-cre mice, the optical fiber was positioned at the cortical surface and the effect of LED inputs of varying intensities on multi-unit neuron spiking was measured throughout the cortical depth. **b.** Grand PSTHs (mean +/− sem across neurons) of the cortical population in conditions with and without the LED at intensities of 8 (left) and 30 mW/mm^2^ (right). The population’s ongoing and sensory evoked activity are unchanged. **c.** The mean change in ongoing firing rate of individual cortical neurons due to LED inputs at 8 mW/mm^2^ (left) and 30 mW/mm^2^ (right), sorted by cortical depth and color-coded according to significance. Significance is determined by the Wilcoxon signed rank test on pre- and post-LED input firing rates across trials. Note that the x-axis range is that shown in Figure S13 for neurons in the NTSR1-cre mice. The spiking of the majority of neurons was unaffected by the LED inputs. **d.** Same as ***c***, but for the sensory response. Note that the x-axis range is that shown in Figure S17 for neurons in the NTSR1-cre mice. The sensory response was unaffected by the LED inputs for all neurons.

**Figure S2.**
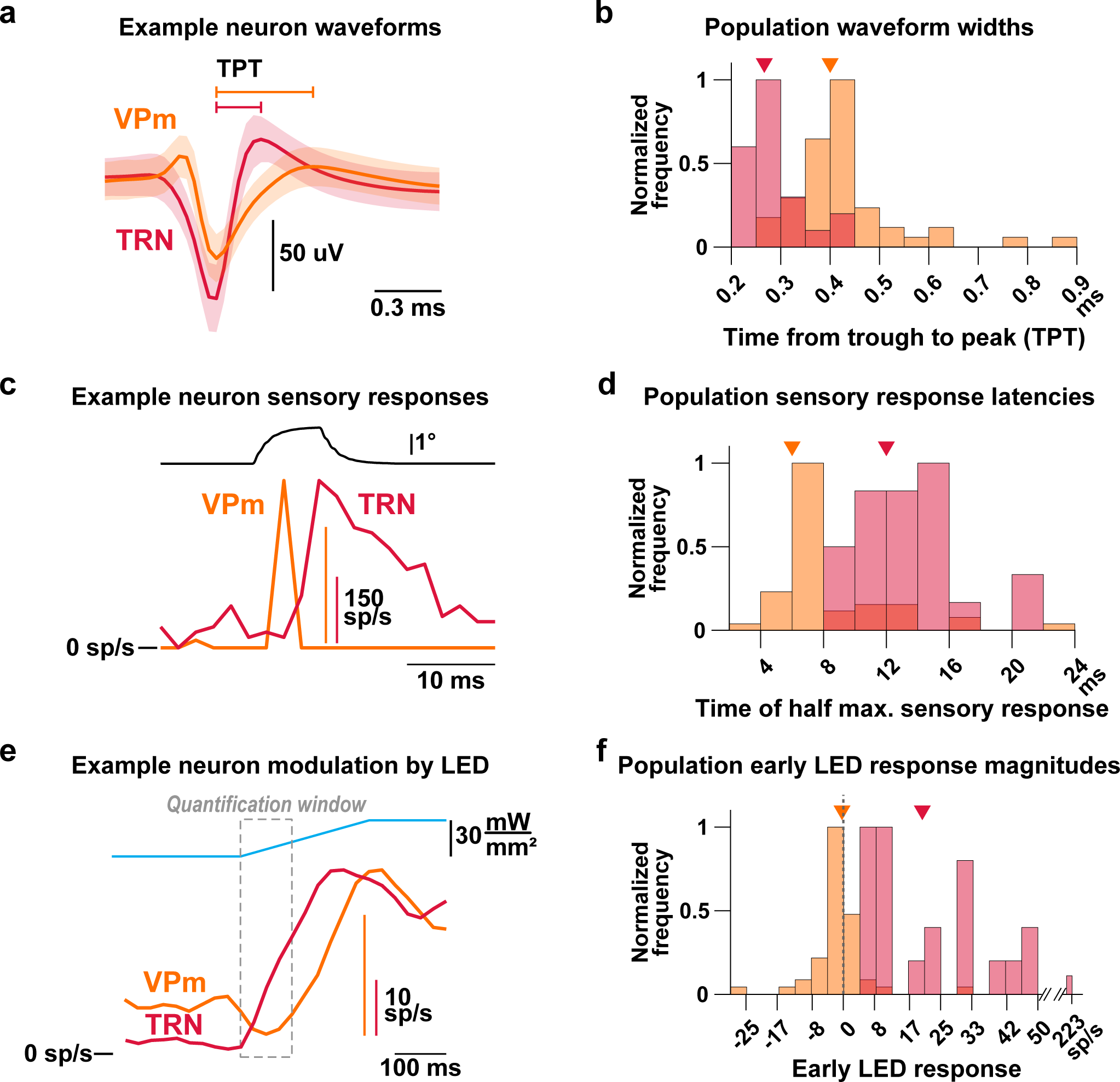
Validated using multiple metrics, identified VPm and TRN neurons are sensory-responsive neurons recorded at the relevant stereotaxic coordinate in NTSR1-cre mice. **a.** Example extracellular spike waveforms of a VPm and TRN neuron (mean +/− standard deviation). The waveform width, quantified by the trough-to-peak time (TPT; see Materials & Methods), of the example VPm neuron is larger than that of the TRN neuron, as observed by others (BarthÓ et al., 2014, Guo et al., 2017, Lewis et al. 2015, Makinson et al., 2017). **b.** The normalized distributions of the TPT of all VPm (orange) and TRN (red) neurons have separable peaks (triangle indicates population median), indicating that TRN neurons have more narrow waveform widths. Excluding VPm neurons with TPTs below 0.333 ms and TRN neurons with TPTs above 0.3 ms did not change any reported results. **c.** Sensory response PSTHs of example VPm and TRN neurons. The TRN neuron’s sensory response is longer lasting in time compared to that of the VPm neuron, as observed by others (Hartings et al., 2003; Ganmor et al., 2010). **d.** The normalized distributions of the time to half of the maximum sensory response of all VPm and TRN neurons have separable peaks (triangle indicates population median), indicating that TRN neurons have longer response latencies. Excluding VPm neurons with latencies above 8 ms did not change any reported results. **e.** PSTHs of the effect of LED activation of L6CT neurons for example VPm and TRN neurons, both whose spiking is enhanced by 30 mW/mm^2^ optogenetic LED input. However, the enhancement of the spiking in the TRN neuron begins sooner than that of the VPm neuron, which is rather transiently suppressed during the first 100 ms of the LED input. **f.** The normalized distributions of the baseline-subtracted magnitude of the LED modulation in the early window outlined in ***e*** of all TRN and VPm neurons are well separated (triangle indicates population median). This indicates that the spiking of all TRN neurons is enhanced during the first 100 ms of the LED input, whereas a large majority of VPm neurons are either suppressed or modestly enhanced. Excluding VPm neurons above a cutoff of 4 sp/s did not change any reported results.

**Figure S3.**
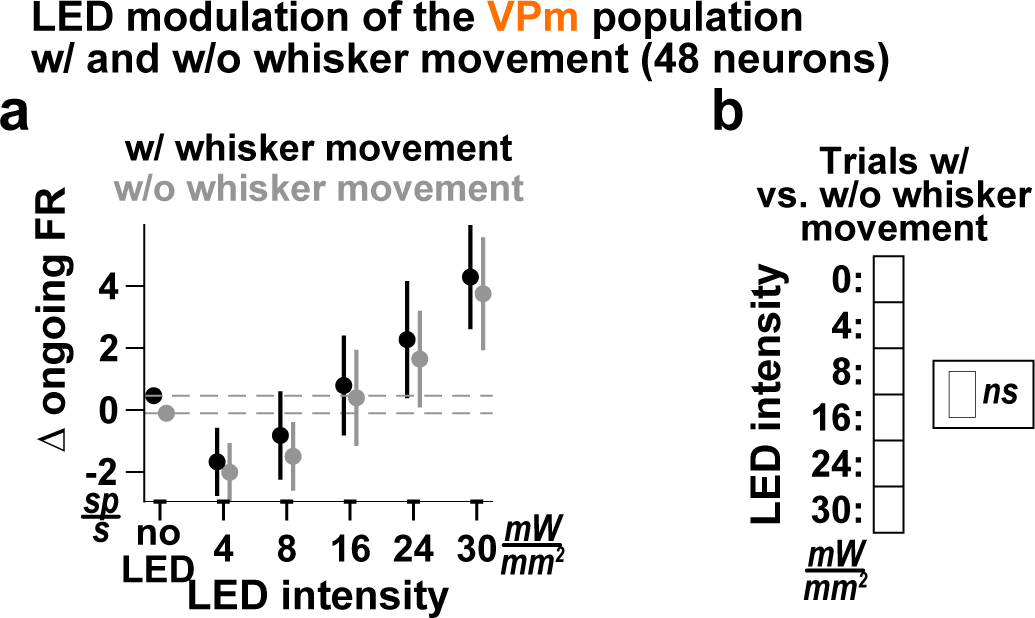
The effect of the LED activation of L6CT neurons on VPm spontaneous activity is nearly identical for trials with and without whisker movement. **a.** The mean +/− sem change in ongoing activity of the VPm population at the various LED intensities in trials with (black) and without (gray) whisker movement. For this analysis, we only included experiments for which there was a minimum of five trials across all LED intensities for both whisker movement conditions. The gray data points thus reflect a subset of the neurons in Figure 2. Note that a more strict requirement on the minimum allowable trials does not change the nature of the results. The color indicates the p-value using the two-sided Wilcoxon signed rank test with a Bonferroni correction. **b.** Confusion matrix indicating the significance level of the change in ongoing firing rate of the VPm population observed in ***a*** when comparing between trials with and without whisker movement at each LED intensity.

**Figure S4.**
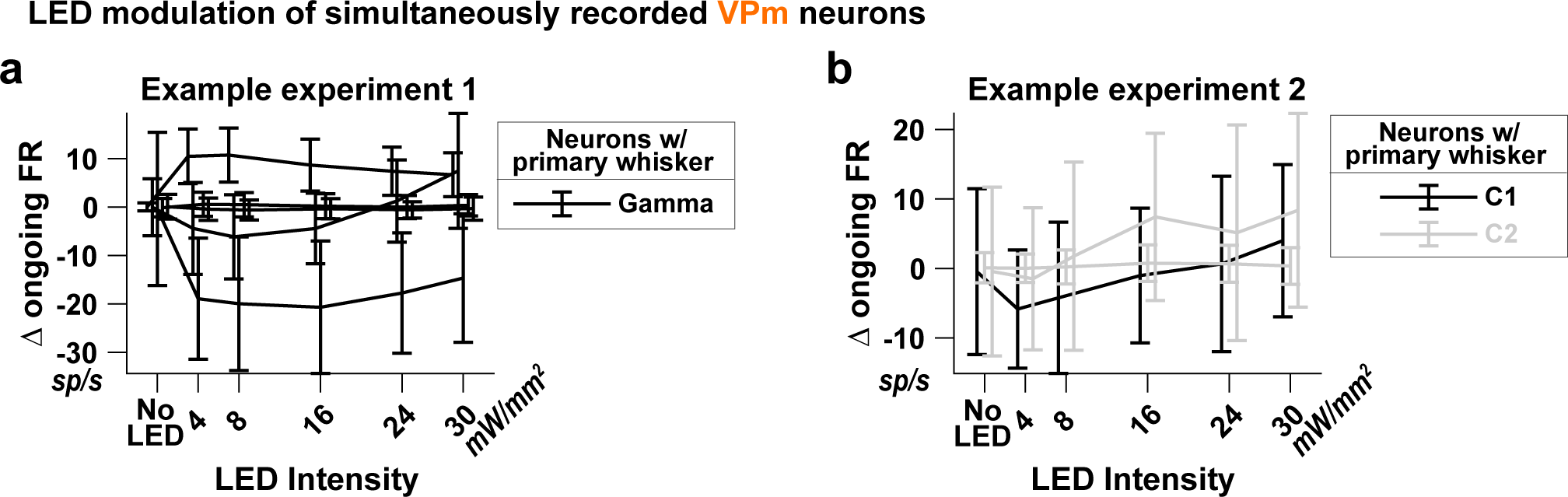
The heterogeneity of effects of the LED inputs on the ongoing activity of VPm neurons is observed across the functional topography of VPm. **a.** Example mean +/− standard deviation change in ongoing firing rate of simultaneously recorded VPm neurons that have the same primary whisker. Therefore, these neurons receive inputs from L6CT neurons of the same functional cortical column. Yet, the LED modulation effect is variable across these neurons, with one enhanced across all LED intensities, one suppressed, two unmodulated, and one bidirectionally modulated neuron. **b.** Same as a, but for an example recording of VPm neurons from two different primary whiskers, thus receiving inputs from L6CT neurons of differing functional cortical columns. Yet, two of the three neurons are bidirectionally modulated. Importantly, these two neurons have differing primary whiskers.

**Figure S5.**
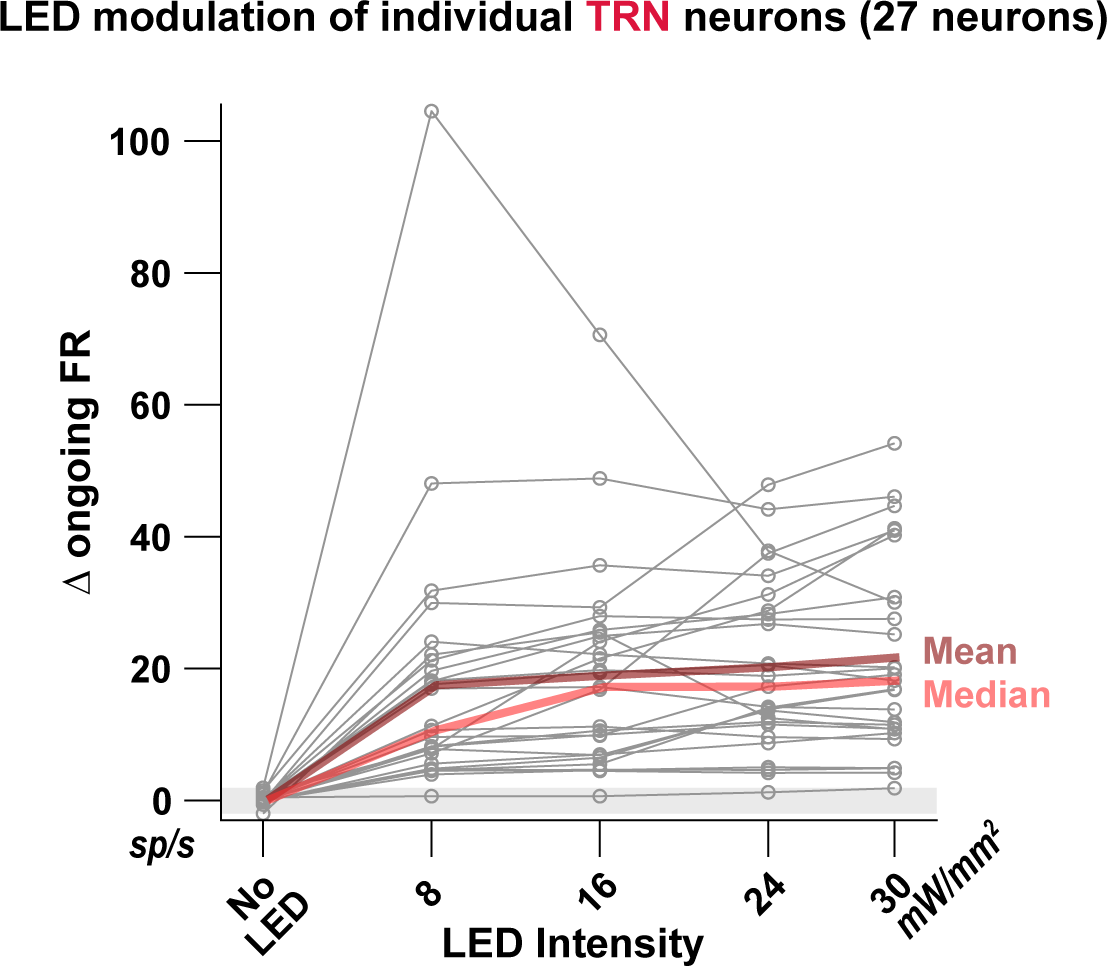
The effect of the LED modulation of L6CT neurons on individual TRN neurons. The mean change in ongoing firing rate of all recorded TRN neurons illustrates relatively homogeneous modulatory effects across the population.

**Figure S6.**
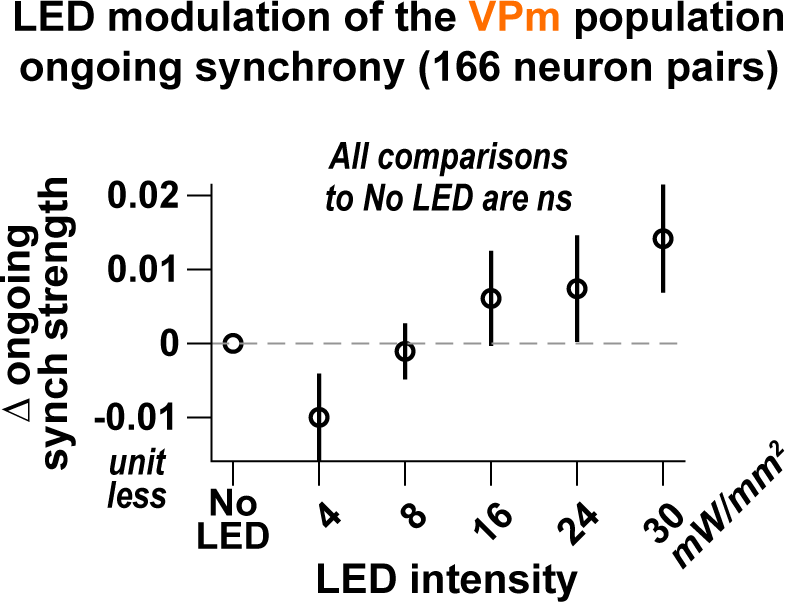
The effect of optogenetic activation of L6CT neurons on the synchrony of ongoing VPm activity is small and nonsignificant. The mean +/− sem change in the VPm population ongoing synchrony strength to ramped LED steps of increasing intensity (statistical comparisons are between the No LED condition and the respective LED-on conditions using the two-sided Wilcoxon signed rank test with a Bonferroni correction).

**Figure S7.**
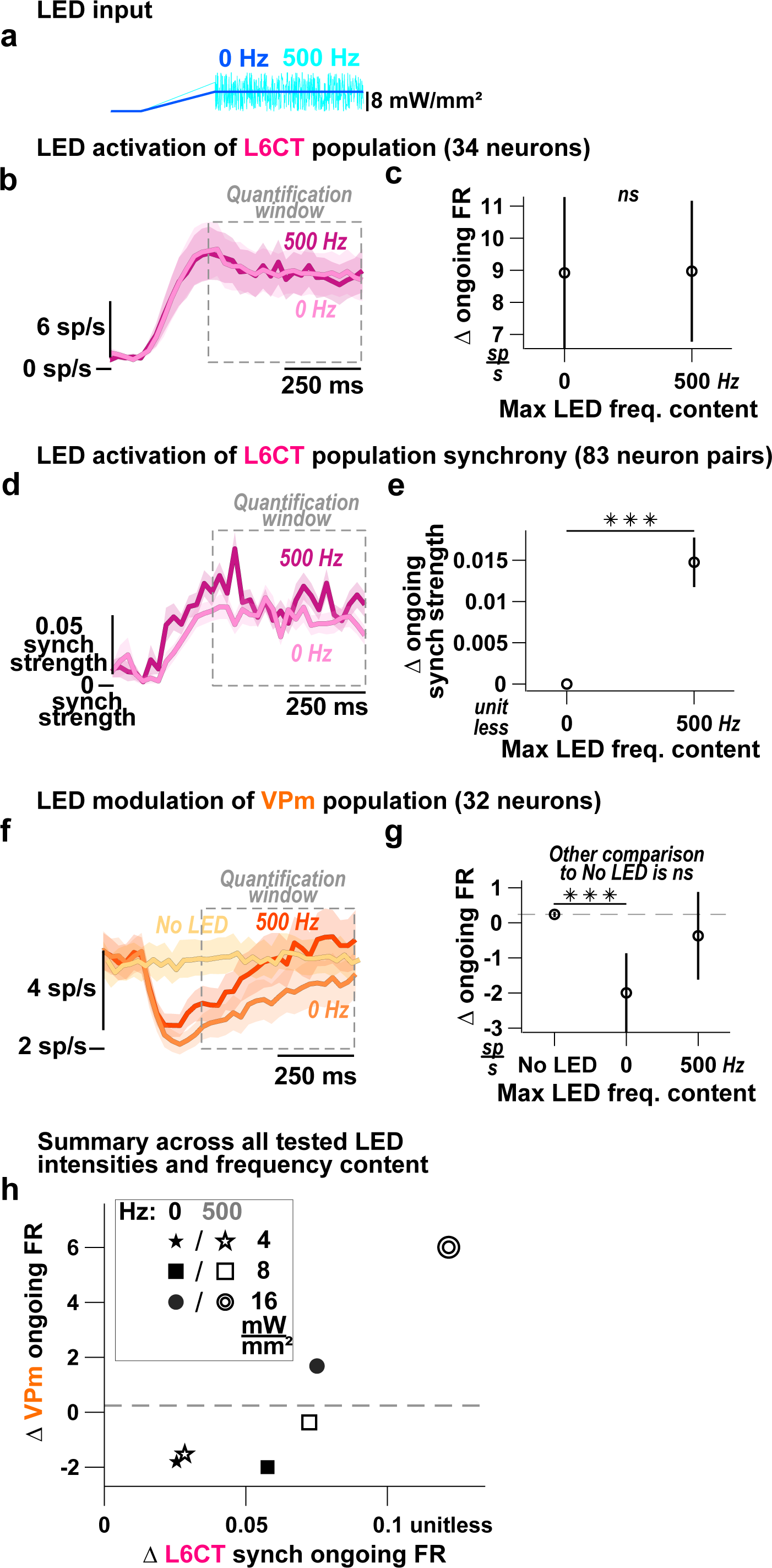
Shifts in L6CT population synchrony across firing rates affect the modulatory effect on VPm activity. **a.** To further probe the effect of L6CT synchrony on VPm ongoing activity, we presented LED inputs with maximum frequency mean of 8 mW/mm^2^ (instead of 16 mW/mm^2^ as in Figure 5). Non-frozen realizations of noise were presented across trials. **b.** Grand PSTHs (mean +/− sem across neurons) of the L6CT population activation by the LED inputs in ***a***. Note that the set of neurons analyzed is not the same as those presented in Figures 4 and 5. Also, this dataset has not been parsed to exclude trials where the mouse moved its whiskers. **c.** The mean +/− sem change in ongoing firing rate of the L6CT population contributing to the PSTHs in ***b*** is unchanged between the 0 and 500 Hz LED inputs (p = 0.12, determined by the two-sided Wilcoxon signed rank test between the conditions). **d.** Grand PSTHs (mean +/− sem across pairs of neurons) of the L6CT population synchrony strength activation by the LED inputs in ***a***. **e.** The mean +/− sem change in synchrony strength of the L6CT population contributing to the PSTHs in ***d*** increases between the 0 and 500 Hz stimuli (p = 8.6e-6, determined by the two-sided Wilcoxon signed rank test between the two conditions). **f.** Grand PSTHs (mean +/− sem across neurons) of the VPm population modulation by the LED inputs in ***a***. **g.** The mean +/− sem change in ongoing firing rate of the VPm population contributing to the PSTHs in ***f*** shifts from a significant decrease in response to the 0 Hz stimulus to no change in response to the 500 Hz LED input (0 Hz: p = 3.8e-4; 500 Hz: p = 6.7e-2; p-value indicates pairwise comparison to No LED condition using the two-sided Wilcoxon signed rank test with a Bonferroni correction). **h.** Summary of the mean effect of L6CT synchrony across firing rates on the mean change in VPm population firing rate. Same marker shape indicates equal LED intensity (either 4, 8, or 16 mW/mm^2^). Filled markers indicate the 0 Hz stimulus and hollow markers indicate the 500 Hz stimulus. Increased L6CT population synchrony with equivalent L6CT population firing rate increases the change in the VPm firing rate across multiple L6CT firing rate pairs, although to varying extents across pairs. Taken together, gradual increases in the synchrony strength of the L6CT population shift the modulatory effect on VPm ongoing firing rates from being suppressive to being enhancive.

**Figure S8.**
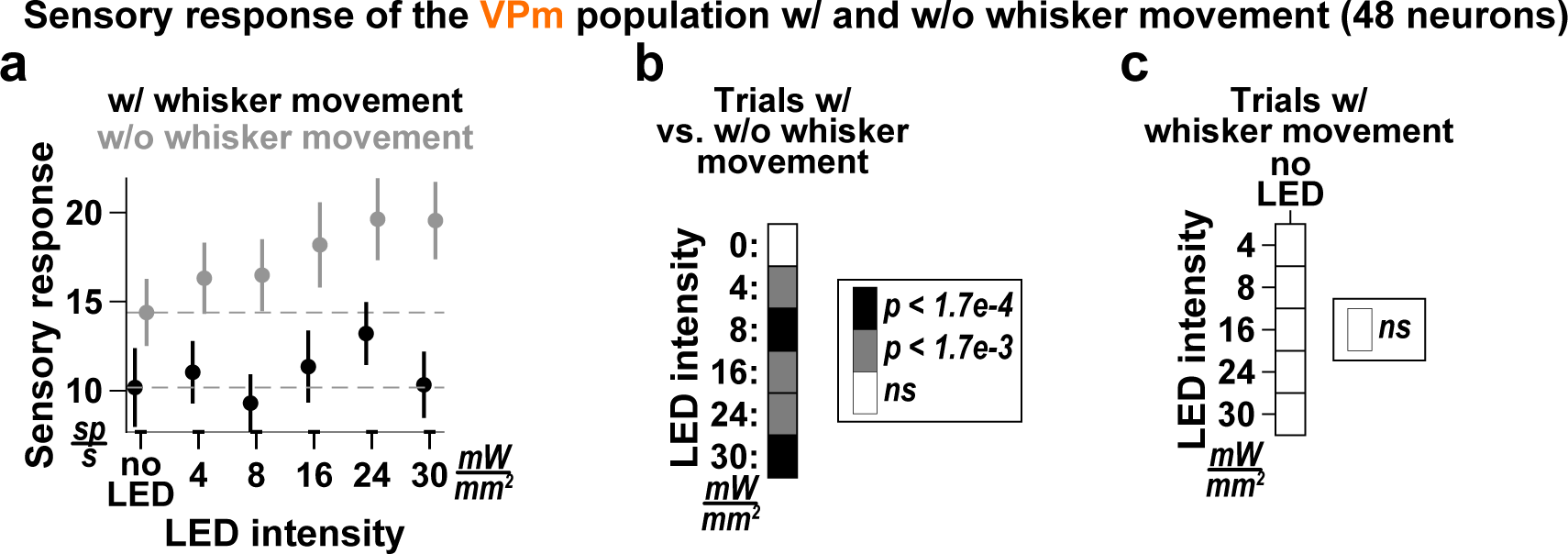
The measured effect of L6CT activation on VPm sensory responses in trials with whisker movement is reduced compared to the effect in trials without whisker movement. **a.** The mean +/− sem sensory response of the VPm population at the various LED intensities in trials with (black) and without (gray) whisker movement. For this analysis, we only included experiments for which there was a minimum of five trials across all LED intensities for both whisker movement conditions. The gray data points thus reflect a subset of the neurons in Figure 7. Note that a more strict requirement on the minimum allowable trials does not change the nature of the results. **b.** Confusion matrix indicating the significance level of the sensory response of the VPm population observed in ***a*** when comparing between trials with and without whisker movement at each LED intensity. The color indicates the p-value using the two-sided Wilcoxon signed rank test with a Bonferroni correction. **c.** Confusion matrix indicating the significance level of the VPm population sensory response between no LED and LED-on conditions observed only in trials with whisker movement; the color indicates the p-value using the two-sided Wilcoxon signed rank test with a Bonferroni correction. No conditions are different from each other.

**Figure S9.**
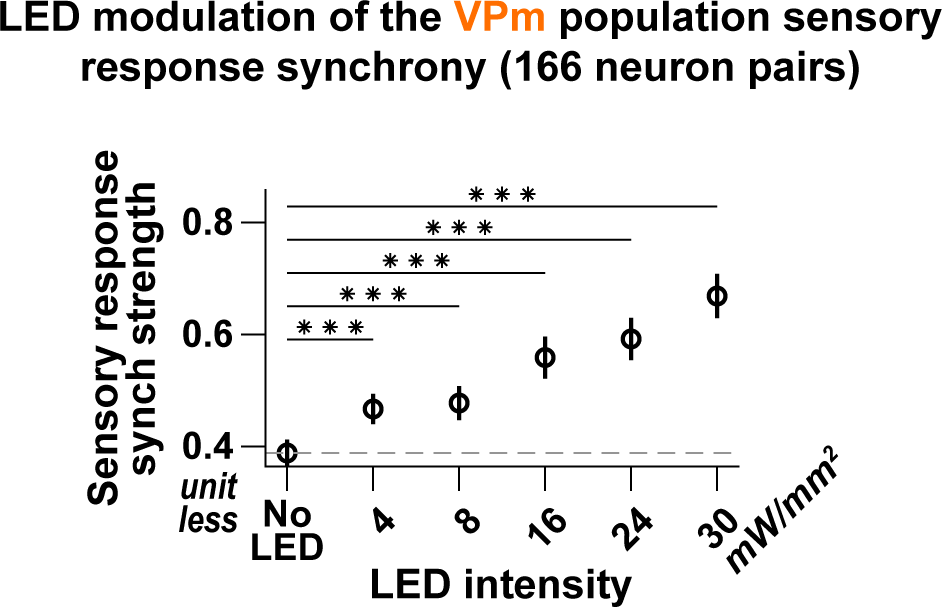
Increasing optogenetic activation of L6CT neurons increases the synchrony of VPm sensory responses. The mean +/− sem VPm population sensory response synchrony strength to ramped LED steps of increasing intensity (statistical comparisons are between the No LED and the respective LED-on conditions using the two-sided Wilcoxon signed rank test with a Bonferroni correction).

**Figure S10.**
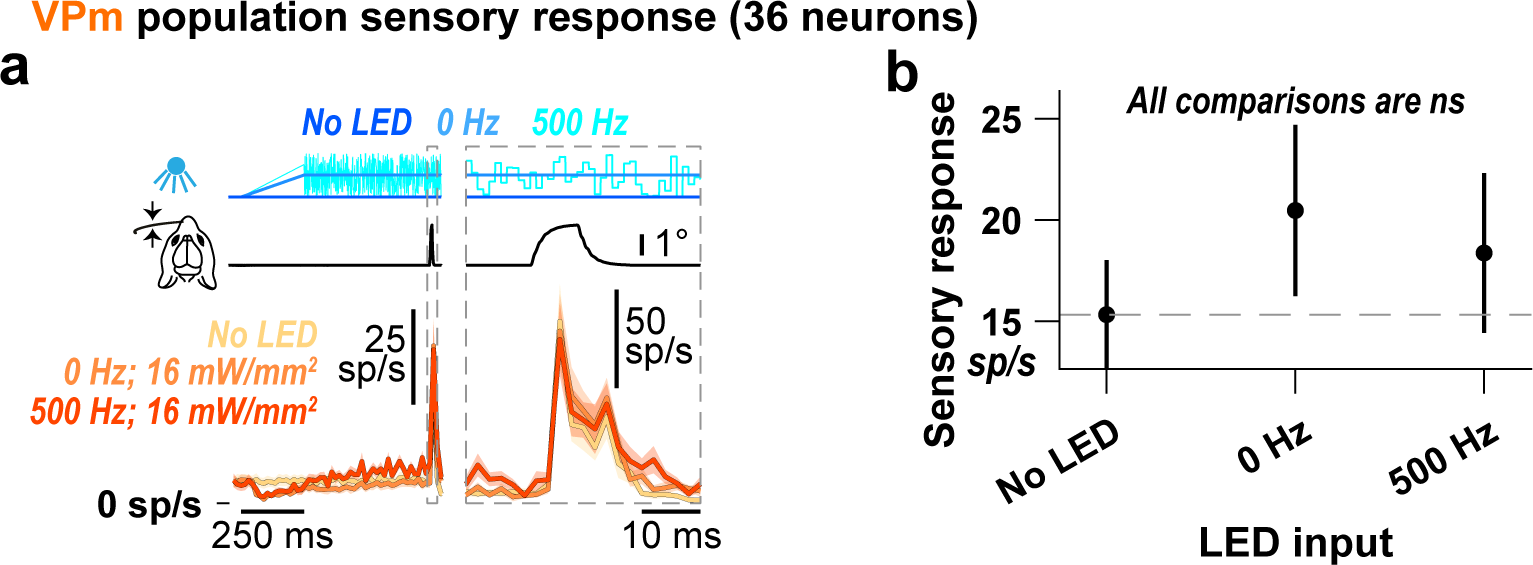
Increased L6CT synchrony does not differentially affect VPm sensory responses. **a.** Grand PSTHs (mean +/− sem across neurons) of the VPm population sensory response with no LED, the 0 Hz LED step, and the 500 Hz frequency modulated LED input, which selectively increases L6CT synchrony and not firing rate when compared to the 0 Hz LED step. Note that this population is a subset of the population presented in Figure 7. The right panel is an expanded view of the same population’s sensory response. Note the change in the vertical scale in the right and left views. **b.** While there is an increasing trend that matches the results presented in Figure 7, the various LED inputs have no significant effect on the baseline-subtracted, stimulus-evoked response of this subset of VPm neurons contributing to the PSTHs in ***a*** (+/− sem). Most relevant, there is no further boost in sensory response between the 0 and 500 Hz conditions (pairwise comparisons are to the No LED or 0 Hz condition using the two-sided Wilcoxon signed rank test with a Bonferroni correction).

**Figure S11.**
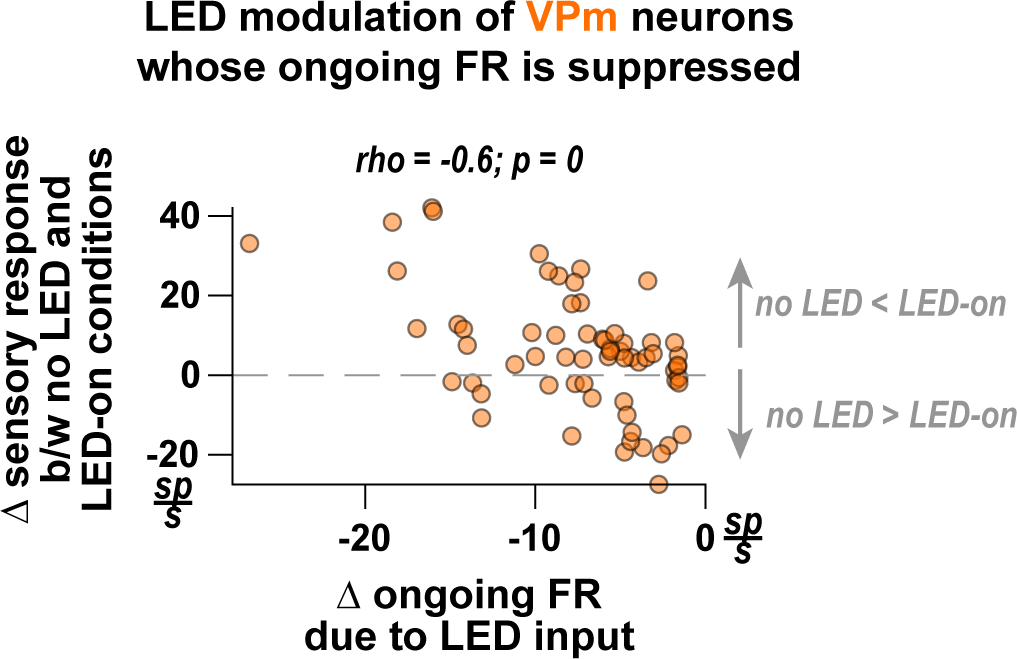
The effect of the LED modulation on the ongoing firing rate of VPm neurons does not trivially predict the effect on the neurons’ sensory response. A simple prediction would be that for neurons whose ongoing firing rate is suppressed by the LED inputs, their sensory response would likewise be reduced. Thus, we present the change in ongoing firing rate of individual VPm neurons due to LED activation of L6CT neurons against the difference in sensory response between no LED and LED-on conditions. Note that we only include neurons’ activity under conditions where the ongoing firing rate is significantly suppressed by LED inputs using the two-sided Wilcoxon signed rank test. Thus, the set of datapoints represent the response of neurons across all tested LED intensities. Under this set, there is a moderate, negative Spearman’s correlation, suggesting a trend where the neurons whose ongoing activity is most suppressed by an LED input rather have a greater sensory response compared to no LED conditions, contrary to the simple prediction. Note that this trend holds even if we only include data under the 4 mW/mm^2^ intensity condition (not shown).

**Figure S12.**
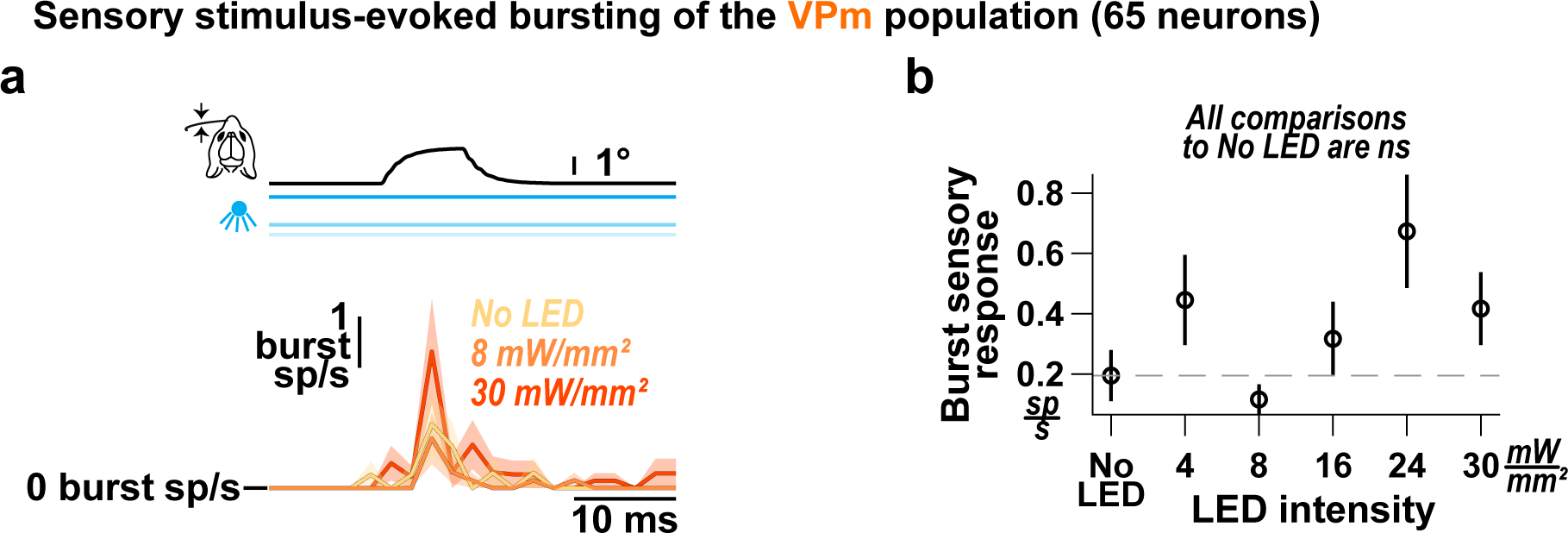
The effect of optogenetic activation of L6CT neurons on VPm stimulus-evoked bursting. **a.** Grand PSTHs of putative T-Type calcium channel burst spikes (mean +/− sem across neurons) of the VPm neuron population. Bursts are defined as two or more spikes with an inter-spike interval less than 4 ms, preceded by 100 ms or more of no spikes (Borden et al., 2022, Lu 1992, Swadlow & Gusev, 2001, Whitmire & Stanley, 2016, Wright et al., 2021). **b.** Optogenetic activation of L6CT neurons has no significant effect on the mean, baseline-subtracted, stimulus-evoked burst response of the VPm population contributing to the PSTHs in ***a*** (+/− sem; No LED: 0.2 +/− 0.08 sp/s; 8 mW/mm^2^: 0.11 +/− 0.05 sp/s, p = 0.27; 30 mW/mm^2^: 0.44 +/− 0.12 sp/s, p = 0.06; p-value indicates pairwise comparison to No LED condition using the two-sided Wilcoxon signed rank test with a Bonferroni correction).

**Figure S13.**
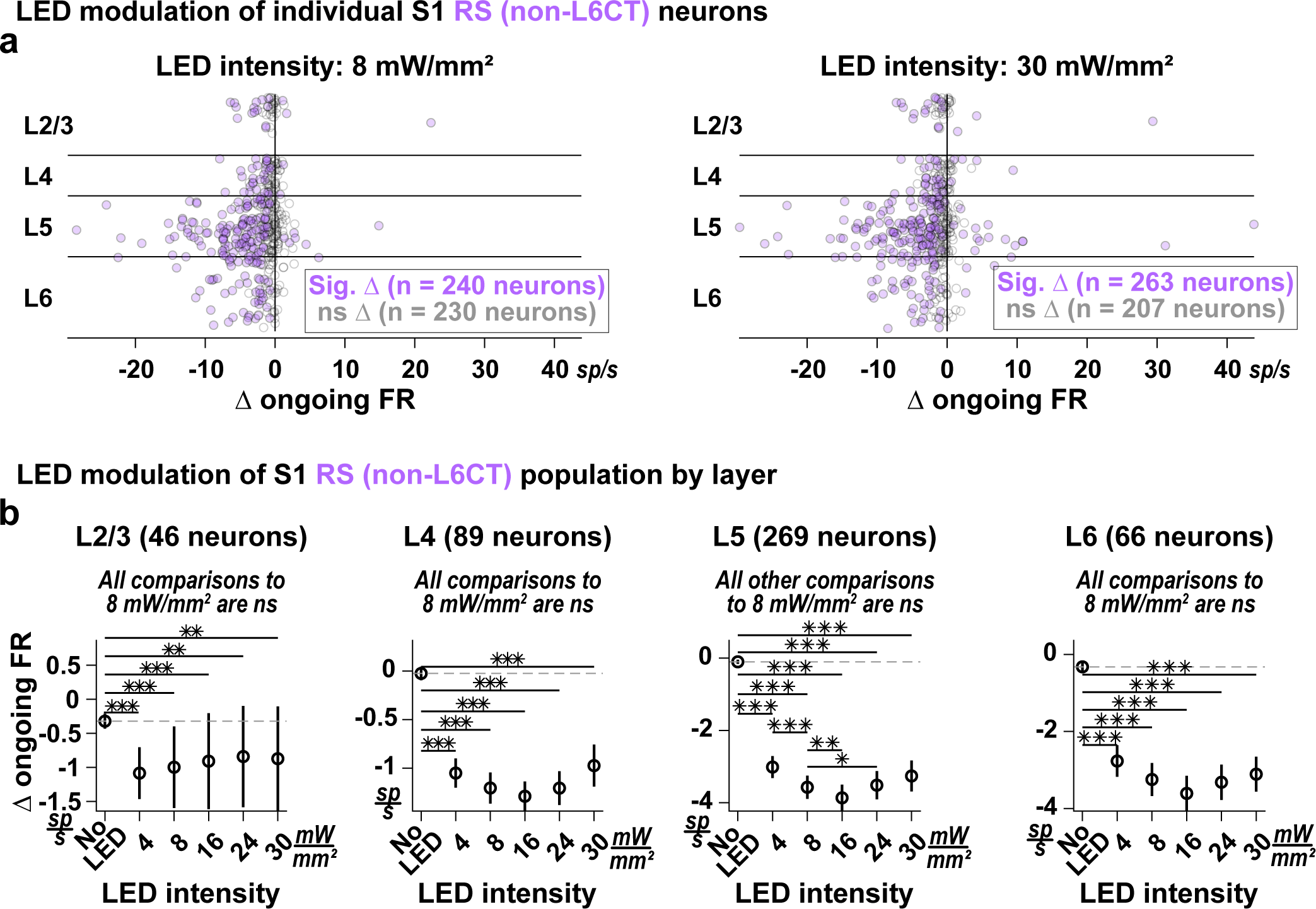
Optogenetic activation of L6CT neurons suppresses ongoing activity in S1 RS neurons across all layers. **a.** The mean change in ongoing firing rate of individual S1 RS (non-L6CT) neurons due to optogenetic activation of L6CT neurons at 8 mW/mm^2^ (left) and 30 mW/mm^2^ (right), sorted by cortical layer and color-coded according to significance (significance is determined by the Wilcoxon signed rank test on pre- and post-LED input firing rates across trials). Across all cortical layers, we observe neurons whose ongoing activity is suppressed or unchanged by optogenetic activation of L6CT neurons, with few neurons that are enhanced. **b.** Optogenetic activation of L6CT neurons decreases the mean +/− sem change in ongoing firing rate of the RS (non-L6CT) neuron population in each cortical layer. Further, when comparing all other LED conditions to the 8 mW/mm^2^ condition where maximal suppression of VPm ongoing activity was observed, although not significant, a small but apparent reversal in the trend emerges in layers 4, 5, and 6 (p-value indicates pairwise comparison to either No LED condition or 8 mW/mm^2^ condition using the two-sided Wilcoxon signed rank test with a Bonferroni correction; * indicates p < 5.6e-3; ** indicates p < 1.1e-3; *** indicates p < 1.1e-4).

**Figure S14.**
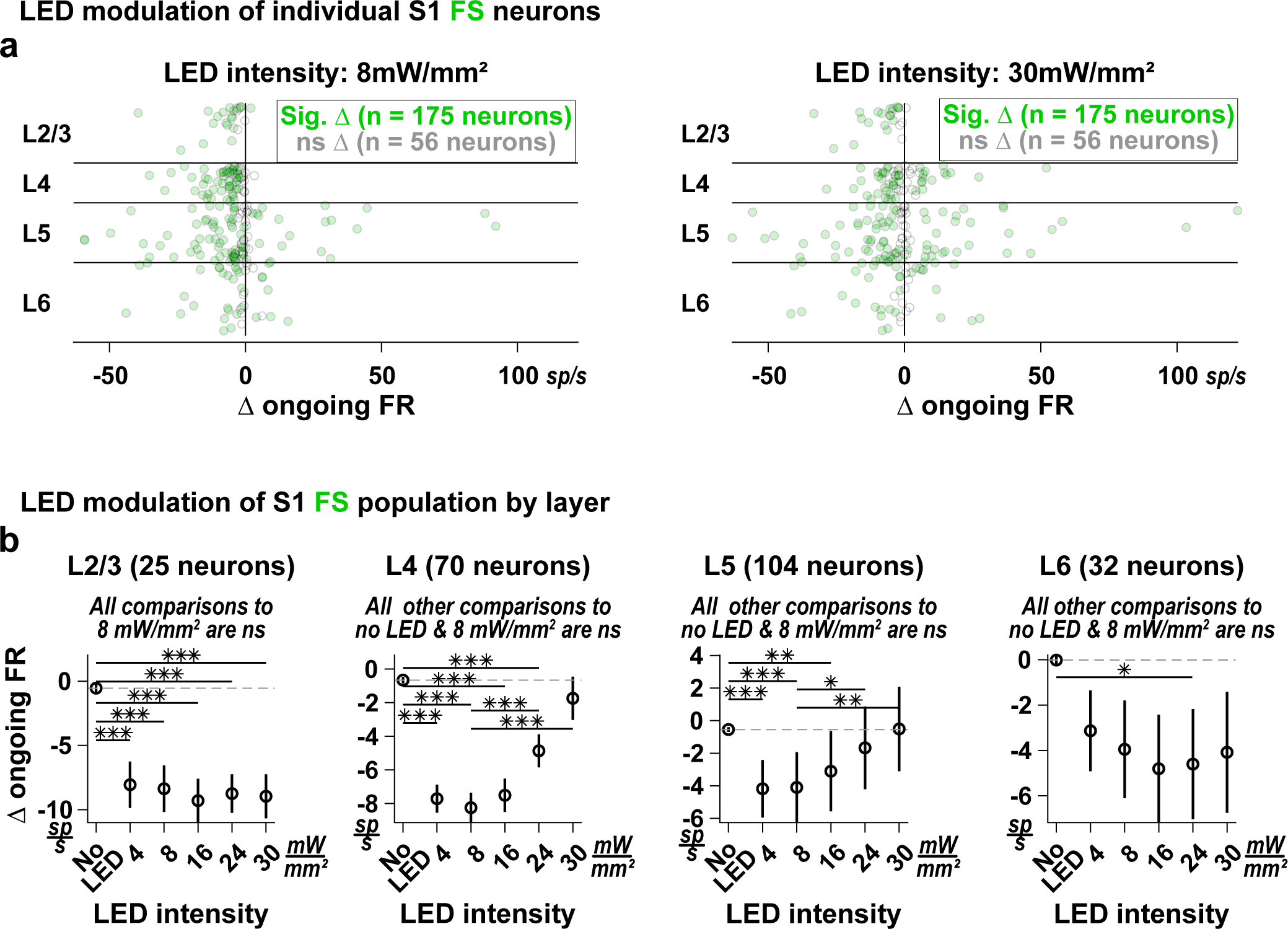
Optogenetic activation of L6CT neurons dynamically suppresses ongoing activity in S1 FS neurons in layers 4 and 5. **a.** The mean change in ongoing firing rate of individual S1 FS neurons due to optogenetic activation of L6CT neurons at 8 mW/mm^2^ (left) and 30 mW/mm^2^ (right), sorted by cortical layer and color-coded according to significance (significance is determined by the two-sided Wilcoxon signed rank test on pre- and post-LED input firing rates across trials). Across all cortical layers, we observe neurons whose ongoing activity is suppressed or unchanged by optogenetic activation of L6CT neurons. With the exception of layer 2/3, all other cortical layers have neuron’s whose ongoing activity is enhanced by optogenetic activation of L6CT neurons, the extent to which depends on the LED intensity. **b.** Optogenetic activation of L6CT neurons decreases the mean +/− sem change in ongoing firing rate of FS populations parsed by cortical layer. Further, when comparing all other LED conditions to the 8 mW/mm^2^ condition where maximal suppression of VPm ongoing activity was observed, a reversal in the trend emerges in layers 4 and 5, such that the effect at 30 mW/mm^2^ is both statistically unchanged from the no LED condition and significantly higher than the effect at 8 mW/mm^2^. Although not significant, a small but apparent reversal also occurs in layer 6 (p-value indicates pairwise comparison to either No LED condition or 8 mW/mm^2^ condition using the two-sided Wilcoxon signed rank test with a Bonferroni correction; * indicates p < 5.6e-3; ** indicates p < 1.1e-3; *** indicates p < 1.1e-4).

**Figure S15.**
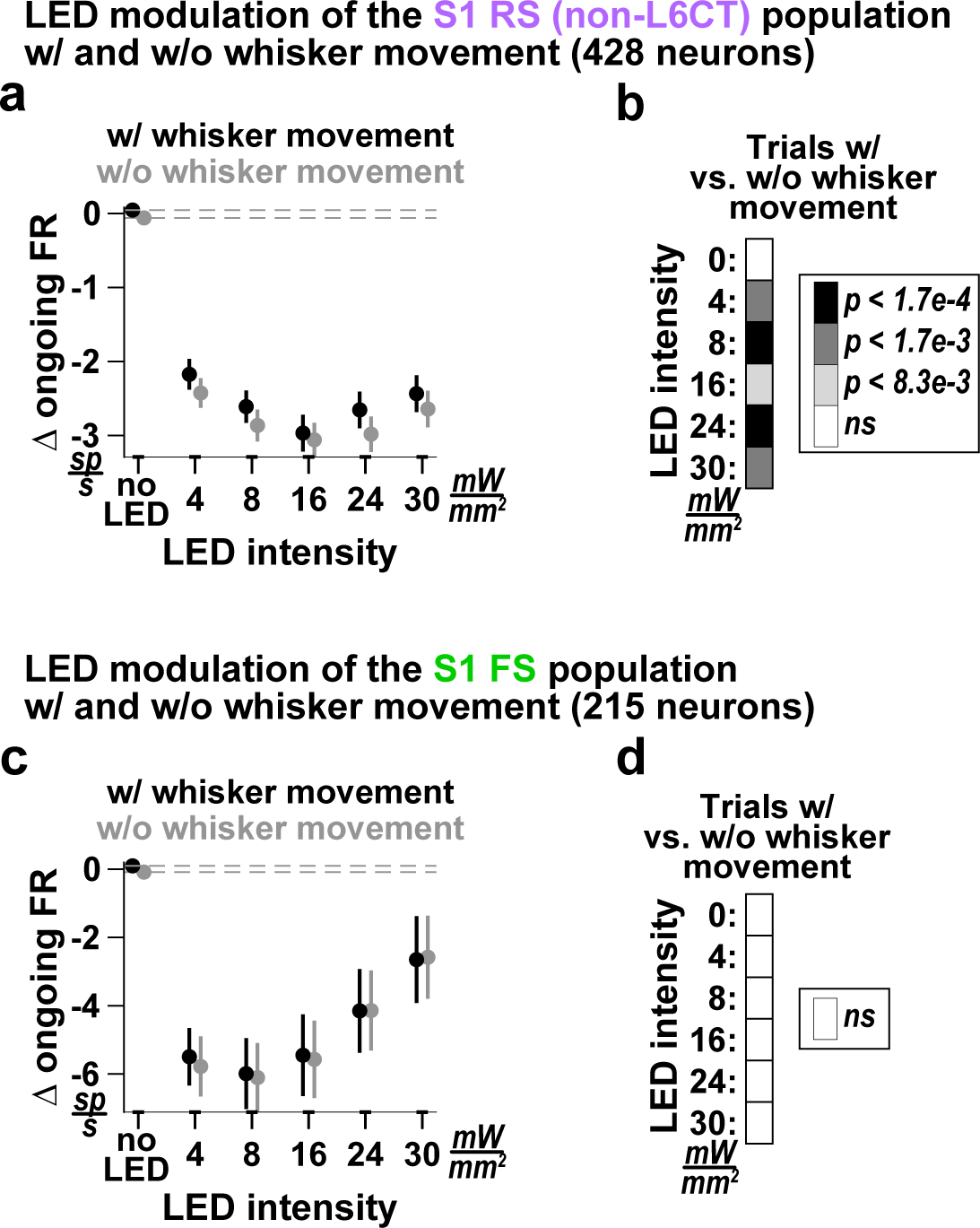
Compared to the effect in trials *without* whisker movement, while the measured effect of L6CT activation on ongoing S1 RS activity is modestly decreased in trials *with* whisker movement, the effect on ongoing S1 FS activity is not associated with any change. **a.** The mean +/− sem change in ongoing activity of the S1 RS (non-L6CT) population at the various LED intensities in trials with (black) and without (gray) whisker movement. For this analysis, we only included experiments for which there was a minimum of five trials across all LED intensities for both whisker movement conditions. The gray data points thus reflect a subset of the neurons in Figure 8. Note that a more strict requirement on the minimum allowable trials does not change the nature of the results. **b.** Confusion matrix indicating the significance level of the change in ongoing firing rate of the RS population observed in ***a*** when comparing between trials with and without whisker movement at each LED intensity. The color indicates the p-value using the two-sided Wilcoxon signed rank test with a Bonferroni correction. **c-d.** Same as ***a-b***, but for the S1 FS population.

**Figure S16.**
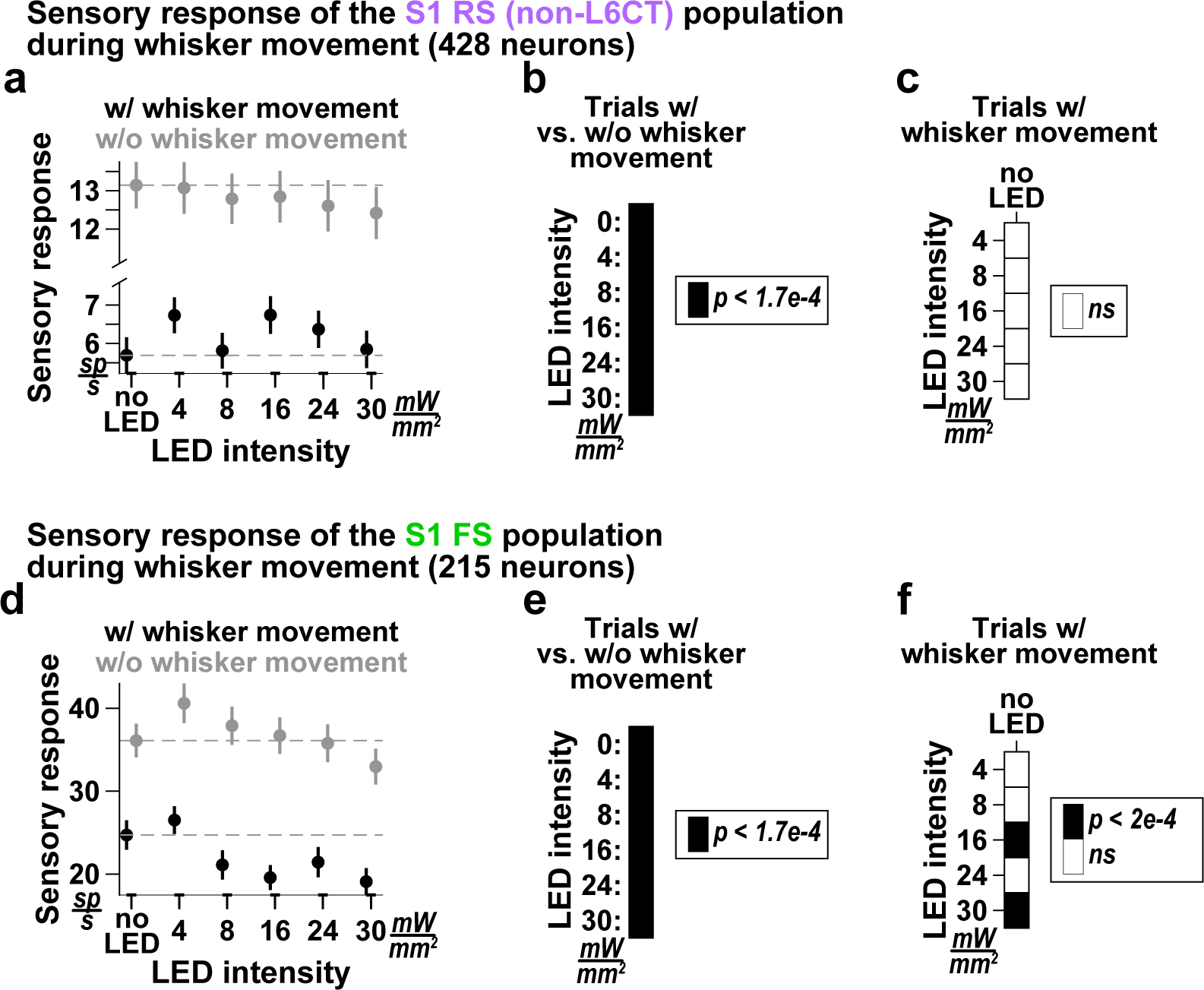
The overall impact of L6CT activation on S1 sensory responses is reduced in trials with whisker movement. **a.** The mean +/− sem sensory response of the S1 RS (non-L6CT) population at the various LED intensities in trials with (black) and without (gray) whisker movement. For this analysis, we only included experiments for which there was a minimum of five trials across all LED intensities for both whisker movement conditions. The gray data points thus reflect a subset of the neurons in Figure 8. Note that a more strict requirement on the minimum allowable trials does not change the nature of the results. **b.** Confusion matrix indicating the significance level of the sensory response of the RS population observed in ***a*** when comparing between trials with and without whisker movement at each LED intensity. The color indicates the p-value using the two-sided Wilcoxon signed rank test with a Bonferroni correction. **c.** Confusion matrix indicating the significance level of the RS population sensory response between no LED and LED-on conditions observed only in trials with whisker movement; the color indicates the p-value using the two-sided Wilcoxon signed rank test with a Bonferroni correction. No conditions are different from each other. **d-f**. Same as ***a-c***, but for the S1 FS population.

**Figure S17.**
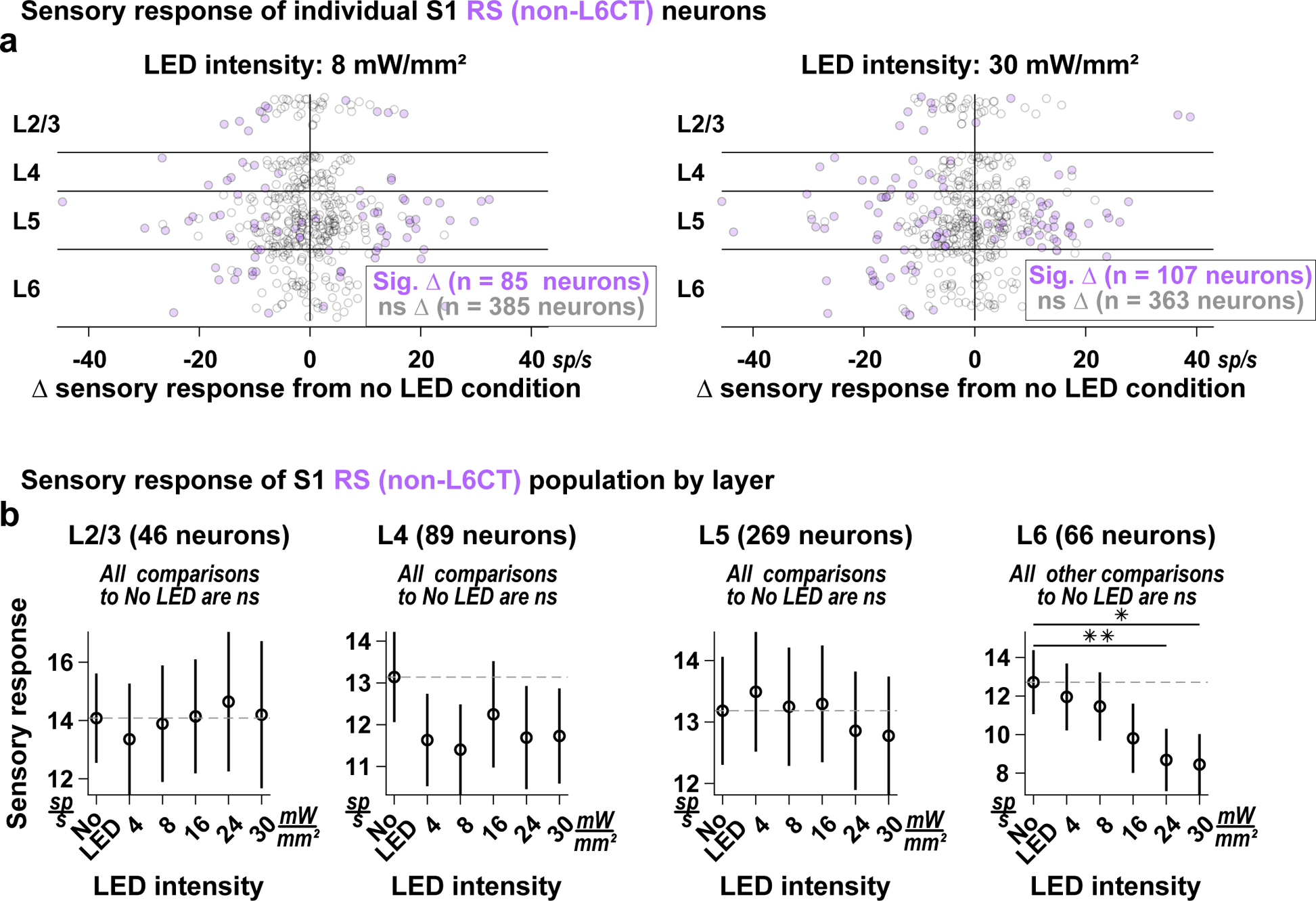
Optogenetic activation of L6CT neurons suppresses S1 RS neurons’ sensory response in cortical layer 6, leaving the sensory response in all other layers unchanged. **a.** The mean change in sensory response from the no LED condition of individual S1 RS (non-L6CT) neurons due to optogenetic activation of L6CT neurons at 8 mW/mm^2^ (left) and 30 mW/mm^2^ (right), sorted by cortical layer and color-coded according to significance (significance is determined by the two-sided Wilcoxon rank sum test on the baseline-subtracted sensory response magnitude across trials in no LED versus LED-on trials). Across all cortical layers, we observe neurons whose sensory response is enhanced, suppressed, and unchanged by optogenetic activation of L6CT neurons. **b.** When parsed by cortical layer, optogenetic activation of L6CT neurons has no effect on the mean +/− sem change in ongoing firing rate of RS (non-L6CT) neuron populations in any layer except layer 6, where sensory responses are suppressed (p-value indicates pairwise comparison to No LED condition using the two-sided Wilcoxon signed rank test with a Bonferroni correction; * indicates p < 1e-2; ** indicates p < 2.0e-3).

**Figure S18.**
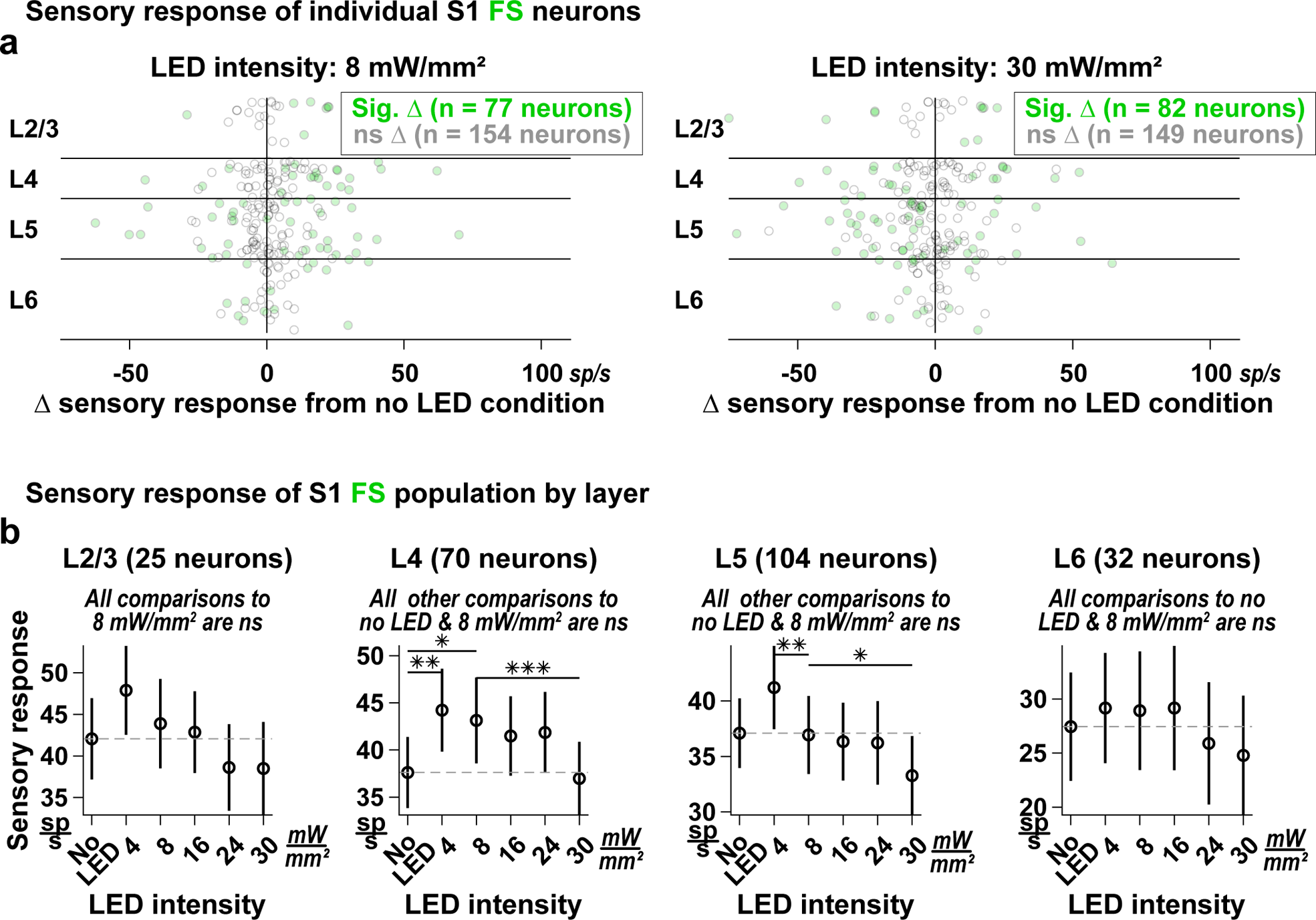
Optogenetic activation of L6CT neurons dynamically modulates the sensory response of S1 FS neurons across all layers. **a.** The mean change in sensory response from the no LED condition of individual S1 FS neurons due to optogenetic activation of L6CT neurons at 8 mW/mm^2^ (left) and 30 mW/mm^2^ (right), sorted by cortical layer and color-coded according to significance (significance is determined by the two-sided Wilcoxon rank sum test on the baseline-subtracted sensory response magnitude across trials in no LED versus LED-on trials). Across all cortical layers, we observe neurons whose sensory response is suppressed, enhanced, or unchanged by optogenetic activation of L6CT neurons. **b.** Optogenetic activation of L6CT neurons dynamically shapes the mean +/− sem sensory response of FS populations parsed by cortical layer, following the general trend of an enhancement of the sensory response at lower LED intensities (although only significant in layers 4 and 5) and a suppression of the sensory response at higher LED intensities. Although the suppression is not significant in any layer, when comparing all other LED conditions to the 8 mW/mm^2^ condition where maximal suppression of VPm ongoing activity was observed, a reversal in the trend of L6CT activation enhancing sensory responses emerges across all layers (p-value indicates pairwise comparison to either No LED condition or 8 mW/mm^2^ condition using the two-sided Wilcoxon signed rank test with a Bonferroni correction; * indicates p < 5.6e-3; ** indicates p < 1.1e-3; *** indicates p < 1.1e-4).

